# A multi-scale fusion CNN model based on adaptive transfer learning for multi-class MI-classification in BCI system

**DOI:** 10.1101/2022.03.17.481909

**Authors:** Arunabha M. Roy

## Abstract

Deep learning-based brain-computer interface (BCI) in motor imagery (MI) has emerged as a powerful method for establishing direct communication between the brain and external electronic devices. However, due to inter-subject variability, inherent complex properties, and low signal-to-noise ratio (SNR) in electroencephalogram (EEG) signal are major challenges that significantly hinders the accuracy of the MI classifier. To overcome this, the present work proposes an efficient transfer learning-based multi-scale feature fused CNN (MSFFCNN) which can capture the distinguishable features of various non-overlapping canonical frequency bands of EEG signals from different convolutional scales for multi-class MI classification. In order to account for inter-subject variability from different subjects, the current work presents 4 different model variants including subject-independent and subject-adaptive classification models considering different adaptation configurations to exploit the full learning capacity of the classifier. Each adaptation configuration has been fine-tuned in an extensively trained pre-trained model and the performance of the classifier has been studied for vast range of learning rates and degrees of adaptation which illustrates the advantages of using an adaptive transfer learning-based model. The model achieves an average classification accuracy of 94.06% (±2.29%) and kappa value of 0.88 outperforming several baseline and current state-of-the-art EEG-based MI classification models with fewer training samples. The present research provides an effective and efficient transfer learning-based end-to-end MI classification framework for designing a high-performance robust MI-BCI system.

## 1. Introduction

Electroencephalogram (EEG) based motor imagery (MI) classification is an critical aspect in brain-computer interfaces (BCIs) which translate brain activities into recognizable machine commands to control the external electronic devices [1, 2, 3, 4, 5, 6, 7]. The BCI allows rehabilitation of neuromotor disorders [8], robotic control [9, 10, 11], speech communication [12, 13], etc. In BCI paradigms, MI classification is the most critical part in which brain signals can be translated into control signals [14, 15]. During MI, desynchronization of neural activities triggers in the primary motor cortex contralateral to the movement which results in a decrease in *μ* − *β* rhythm known as event-related desynchronization (ERD) [16, 17]. It is usually followed by an increase in the *β* rhythm called event-related synchronization (ERS) when MI ceases to exist. Thus, the main goal is to classify different MI tasks according to ERS and ERD characterized by changing of the power spectrum of *μ* (8–14 Hz) and *β* (14–30 Hz) bands.

In this regard, common spatial pattern (CSP) [18] and filter bank CSP (FBCSP) [19] are efficient methods in extracting discriminative features that represent ERD and ERS for MI classification. FBCSP finds a set of linear projections that maximize the differences in the variance of the MI classes by employing filtered signals from various frequency bands which have shown to be effective in improving accuracy.

However, brain signal from EEG has various distinct characteristics (i.e., non-linearity, uniqueness, and non-stationary behaviors) which significantly vary with the human brain and depend on the mental state of the individual subjects [20]. Moreover, due to the presence of noise from various muscle artifacts, fatigue, change of environment, and internal body states may significantly alter the characteristic of EEG signals [14, 15]. Hence, aforementioned factors make it challenging to improve the signal-to-noise ratio (SNR) for better performance in MI classification.

With the recent advancement of deep learning (DL), it demonstrates superior performance in various applications [21, 22, 23] In this regard, convolutional neural network (CNN) has inherent capability of adapting non-linear EEG signal and extracting important feature information automatically in MI-BCI classification [24, 25]. Thus, several studies are geared towards EEG signal classification employing CNN [24, 25, 26, 27, 28, 29, 30]. More recently, there are various CNN based methods [31, 32, 33, 34, 35] have been developed which demonstrate excellent performance in EEG MI classification.

However, DL-based algorithms contain large numbers of trainable model parameters which require a significant amount of training data and lead to increase in computation time to train the classifier [36, 37]. In this regard, transfer learning (TL) based approach is an effective strategy utilizing pre-trained weights from different subject cases [37, 38, 39, 40, 41]. In a recent development, TL method have been employed DL based in the system, in which, a FBCSP based EEG representation utilizing knowledge distillation techniques has been used for MI-BCI classification with a fine-tuned CNN model [36, 42]. Additionally, a subject-independent deep CNN model has been developed using spectral-spatial input generation for MI-BCI system [37]. A hybrid deep neural network utilizing transfer learning has been applied for multi-class MI decoding for better performance [41]. Furthermore, adaptive transfer learning-based deep CNN has been employed for EEG MI classification which has demonstrated significant improvement in classification accuracy compared to subject-dependent model [40].

Although, the CNN-based models have achieved better results for EEG-based MI classification, there are various issues which have caused to hinder the performance of the classifier. Firstly, current CNN-based models only consider a single convolution scale for extracting features from MI EEG signal. Such a strategy may not be suitable to capture distinguishable features of various non-overlapping canonical frequency bands of EEG signals efficiently [43]. Secondly, an important discriminative feature extraction accounting for event-related desynchronization and synchronization (ERD/ERS) from MI EEG signal has been often ignored which limits classifier ability to learn important semantic features from the raw EEG data. Thirdly, a handful of research has been geared towards establishing the efficient transfer learning framework to address the challenge of inter-subject variability between different subjects in the BCI system which requires fine-tuning the model during target MI subject classification [44]. Furthermore, feature extraction from multiscale CNN considering adaptation-based transfer learning yet to be designed for full integration into end-to-end DL workflow which is the main bottleneck for the deployment of robust BCI applications with good classification accuracy. Therefore, the goal of the current work is to develop a robust deep learning framework accounting for inter-subject variability of different subjects and further improve the performance of the model by employing the subject-adaptive transfer learning strategy to achieve better accuracy with fewer training samples.

Motivated by the aforementioned challenges and shortcomings, the current work proposes a transfer learning-based multi-scale feature fused CNN (MSFFCNN) for EEG-based multi-class MI classification. The major contributions and findings of the present research work can be summarized as follows.

- The current work designs an efficient multiscale CNN (MSCNN) architecture to capture the semantic features of EEG signals from multiple convolutional scales for four distinguishable frequency bands *α,β, δ*, and *θ* from raw EEG signal to enhance the performance of MI classifier.
- The present study designs a new FBCSP with the one-vs-rest (OVR) CNN block (OVR-FBCSP CNN) for extracting the discriminative spatiotemporal CSP features of eventrelated desynchronization and synchronization (ERD/ERS) suitable for multiclass MI classification tasks.
- In the current work, 4 different model variants of MSCNN including subject-specific, subject-independent, and subject-adaptive classification model considering two different adaptation configurations have been proposed to exploit the full learning capacity of the classifier.
- The performance of two different subject-adaptive models have been extensively studied for vast range of learning rates and degree of adaption to explore the adaptation capability of the proposed model accounting for inter-subject variability from different subjects.
- Current study illustrates the advantages of using adaptive transfer learning-based model over the subject-specific and subject-independent models in terms of the overall performance of the classifier by achieving the best average classification accuracy outperforming several state-of-the-art models with fewer training samples.

The proposed framework requires less training data and computation time suitable for designing efficient and robust real-time human-robot interaction. The current study effectively addresses the shortcoming of existing CNN-based EEG-MI classification models and significantly improves the classification accuracy. The paper is organized as follows: Section 2 describes the dataset and proposed EEG data representation; OVR-FBCSP have been described in section 3; section 4 introduces the proposed MSFFCNN framework; section 5 describes the transfer learning methods; sections 6 and 7 deal with the relevant finding and discussion of the proposed classifier. Finally, the conclusions and prospects of the current work have been discussed in section 8.

## 2. Dataset and EEG input data representation

### 2.1 Dataset

For the current study, the performance of the MSFFNN model has been evaluated on the BCI competition IV-2a dataset which contains EEG data from 9 subjects for 4 different MI classes (i.e., left hand, right hand, feet, and tongue) [45]. The recorded EEG data comprises two sessions from each subject with the first and second session consisting of training and test data, respectively. Each session was recorded for 288 trials (72 trials per class) from 22 EEG channels with a sampling frequency of 250 Hz according to standardized international 10-20 electrode system as shown in Fig. 1-(a). In a single trial, there was a cue followed by 4 *sec* of MI activity from each subject for each of the four classes as illustrated in Fig. 1 -(b, c).

**Figure 1:**
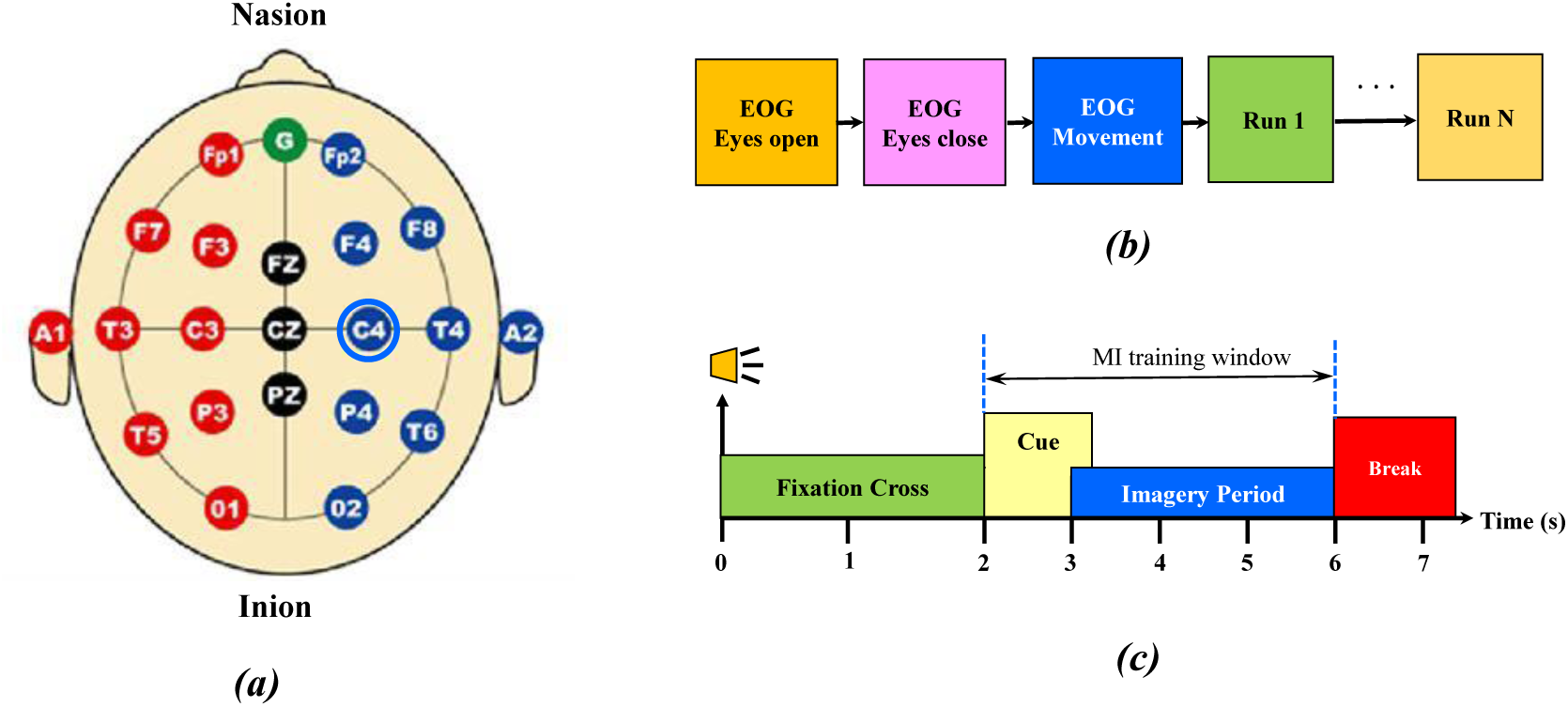
(a) Schematics of EEG electrodes positioning in international 10-20 standardized electrode system; (b) timing scheme of each session; (c) timing scheme of each trial with 4 seconds of MI activity.

### 2.2 Proposed EEG data representation

Since DL methods in MI-BCI systems requires relatively large numbers of EEG training data to achieve good classification accuracy, current work proposed a 1D EEG segment as input data representation. Generally, in MI-BCI systems, inputted data are considered as 2D array combining spatial information of muti-channels (electrodes) and corresponding time-series data for each trial [40]. Such representation ignores the positional electrode distribution in the actual acquisition device and may lead to complex multi-channel correlations instead of simple adjacent relationships [25]. To circumvent such issues, 1D EEG segment from each electrode of fix time window of 4 *sec* has been considered for a particular subject to segment the signals related to MI task as illustrated in Fig. 2 In the proposed method, input data from each channel can be represented by 1D vector 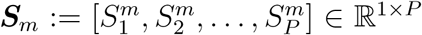 where *m* is the number of channels; *P* = 1000 is the number of sampling points for a single MI task. Such simple and effective data representation results in a substantial increase in the number of training samples containing only time-varying information and time dependence of the signals related to the MI activity. The proposed method can effectively illuminate unnecessary features such as the channel-related spatial information and correlations between electrodes [25, 40].

**Figure 2:**
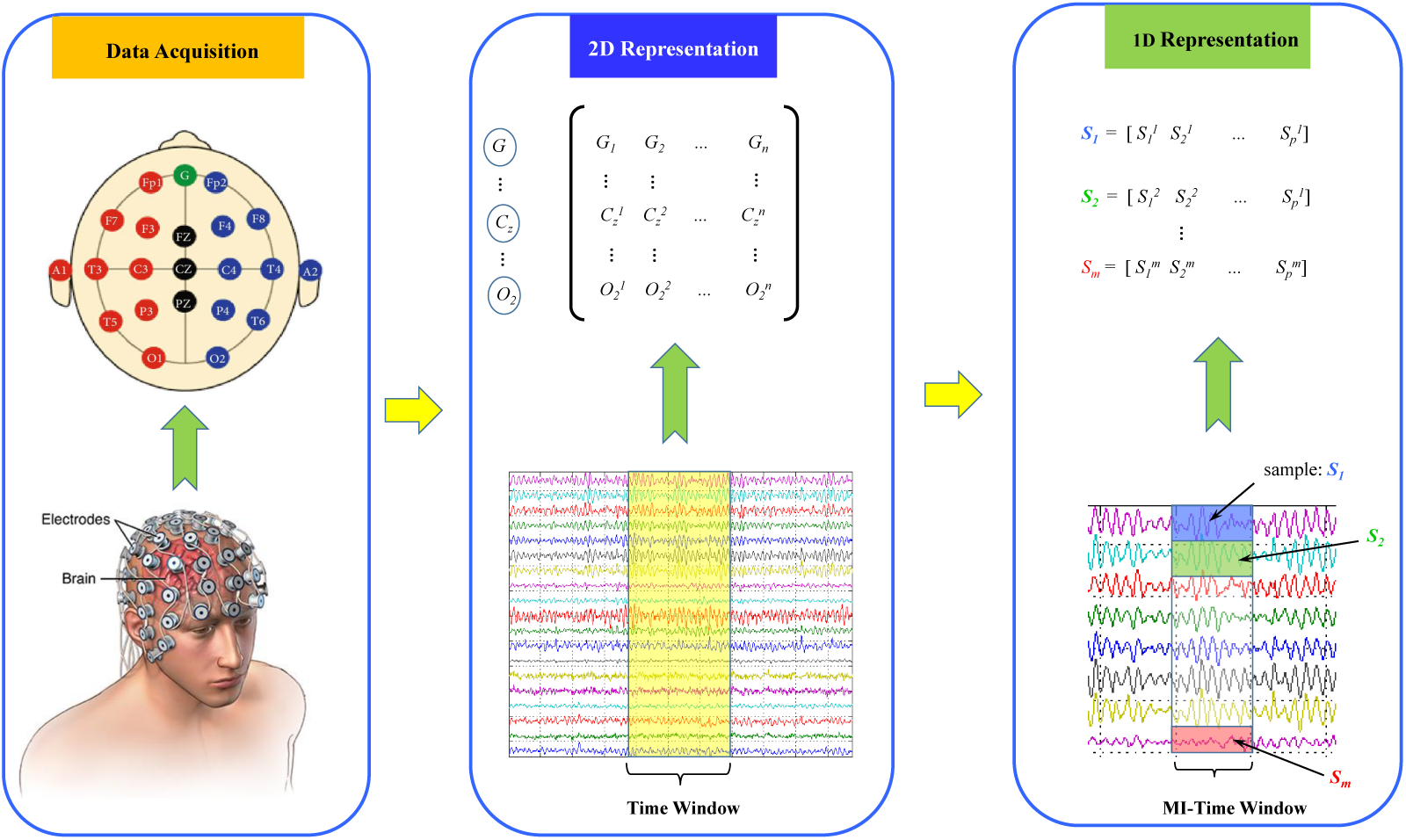
Illustration of proposed EEG input data representation where acquired 2D data from the electrode distribution has been segmented into 1D vector during MI-activity time window.

## 3. One-vs-Rest Filter Bank Common Spatial Pattern

In the present work, FBCSP with the one-vs-rest (OVR) method has been utilized for extracting the spatiotemporal-frequency features of event-related desynchronization and synchronization (ERD/ERS) [46]. The OVR-FBCSP network consists of several FBCSP blocks where the segment of EEG signal can be decomposed through a filter bank that contains an array of multiple types II Chebyshev sub-bandpass filters to extract the discriminative CSP features, for multi-class MI-BCI system [34, 41, 47].

### 3.1. Spatial feature extraction

By employing OVR-CSP, spatial filtering has been performed by linearly transforming EEG signal to obtain feature information using

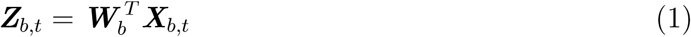

where *t* is numbers of EEG sample per channel; *T* denotes transpose operation; ***X***_*b,t*_ ∈ ℝ^1*×t*^ represents the single-trial EEG segment from *b*-th band-pass filter of *t*-th trial; ***Z***_*b,t*_ ∈ ℝ^1*×t*^ denotes OVR-FBCSP features; 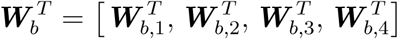 is the weight of OVR-FBCSP filter in which 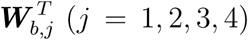 represents CSP projection matrix. Following eigenvalue decomposition problem [46], the transformation matrix 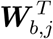 can be obtained to yield optimal discriminating feature variances for multi-class MI as

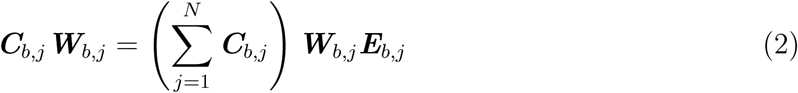

where *N* is a number of MI classes; ***C***_*b,j*_ is the covariance matrix of *b*-th band-pass filtered EEG signal of respective *j*th MI class; ***E***_*b,j*_ is the diagonal matrix containing eigenvalues of ***C***_*b,j*_. Utilizing ***W***_*b,j*_ from Eq. 2, spatial filtered signal ***Z***_*b,t*_ maximizes the differences in the variance of MI classes. For each class, *m* pairs of CSP features of the *t*^*th*^ time window for *b*th band-pass filtered EEG signal can be expressed as:

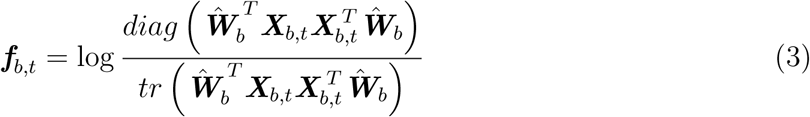

where ***f***_*b,t*_ ∈ ℝ^2*m*^ is the OVR-FBCSP output; ***Ŵ***_*b*_ denotes the first *m* and last *m* columns of ***W***_*b,j*_ (*j* = 1, 2, 3, 4); *diag*(·) is diagonal elements of the square matrix; *tr*(·) is the trace of the matrix. Note that *m* = 2 has been used for BCI IV 2a dataset. The OVR-FBCSP feature vector corresponds to *t*^*th*^ time window can be expressed as

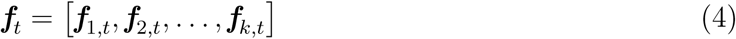

where ***f***_*t*_ ∈ ℝ^1*×*2*mk*^ (*t* = 1, 2, …, *n*); *n* total numbers of time windows (i.e., trials); *k* is the total number of band-pass filters. Training data that comprises extracted feature 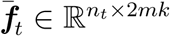 and corresponding true class labels 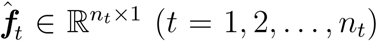; *n*_*t*_ total numbers of trials in training data to make a distinction from evaluation data. Finally, the output matrix from OVR-FBCSP for each 1s time windows each has been attached to CNN layer called OVR-FBCSP CNN for spatial feature extraction (see Section 4.3).

### 3.2 Signal preprocessing

Each MI trial of 4 *sec* long EEG signal has been segmented into 4 parts of 1 *sec* time windows and fed into OVR-FBCSP to obtain spatial features as illustrated in Fig. 6. In OVR-FBCSP, the inputted EEG signal data have been passed through a filter bank that contains an array of a total of 12 bandpass filters including 2-6, 4-8, …, 24-28 Hz. Each filter with a bandwidth of 4 Hz and an overlap of 2Hz has been employed covering the frequency range 2-32Hz.

## 4. Proposed multi-scale feature fused CNN (MSFFCNN)

Achieving good accuracy in MI-BCI subject classification can be challenging due to the high variability and non-stationary characteristic of the raw EEG data. In this regard, CNN has demonstrated the effectiveness of extracting relevant features. Additionally, it is efficient in learning the hierarchical representations of complex high dimensional EEG data. CNN is a feed-forward artificial neural network generally consisting of alternating convolutional and subsampling layers to extract important local features information between adjacent elements of the feature vector, and a fully connected (FC) layer at the end for final classification. Present work proposed an efficient end-to-end MSFFCNN framework in order to improve the accuracy and performance of multi-class MI classification tasks. The MSFFCNN model consists of multi-scale CNN (MSCNN), One-vs-Rest Filter Bank Common Spatial Pattern CNN (OVER-FBCSP CNN), Conv-pool layer, and fully connected layers as illustrated in Fig. 3. Each network block and the corresponding algorithm have been detailed in the subsequent sections.

**Figure 3:**
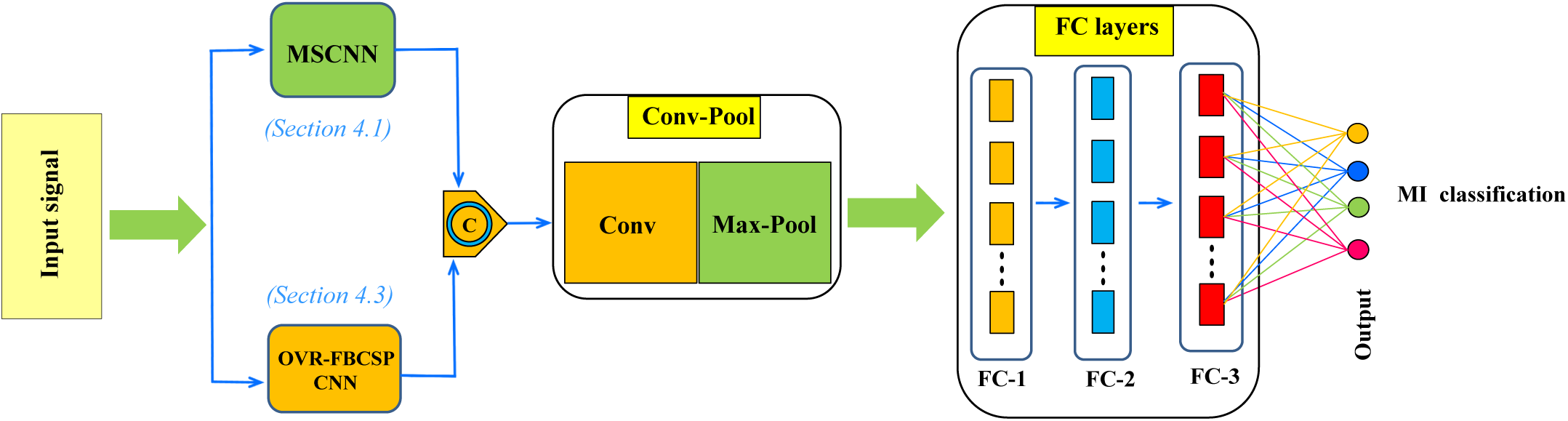
The overall network architecture of proposed end-to-end MSFFCNN consists of (a) multi-scale CNN (MSCNN), (b) One-vs-Rest Filter Bank Common Spatial Pattern CNN (OVER-FBCSP CNN) (c) conv-pool layer (d) fully connected (FC) layers for multiclass MI classification.

### 4.1 Multi-scale CNN block (MSCNN)

In Inputted EEG data, there exist multiple non-overlapping canonical frequency bands where each band signify distinct behavioral states [48, 49]. For MI-BCI systems, *α* (8-13 HZ)and *β* (13-30) bands are the most important [50], since the increase/decrease of the power spectrum of these bands result in ERS/ERD, respectively [51, 52]. Whereas, low frequency *δ*-bands (2-4 Hz) carry important class-related information [14, 53, 54]. Additionally, *θ*-band (4-8 Hz) differs during the left/right-hand MI which is helpful during MI-BCI classification process [33, 55, 56]. Hence, these four non-overlapping bands (i.e., *α, β, δ*, and *θ*) have been considered for feature extraction by employing filter bank of 2-4 Hz, 4-8 Hz, 8-13 Hz, and 13-30 Hz in corresponding frequency bands. In the present study, a multi-scale CNN (MSCNN) with multi-scale convolution blocks (MCBs) has been proposed to extract feature information from each canonical frequency band efficiently as shown in Fig. 4.

**Figure 4:**
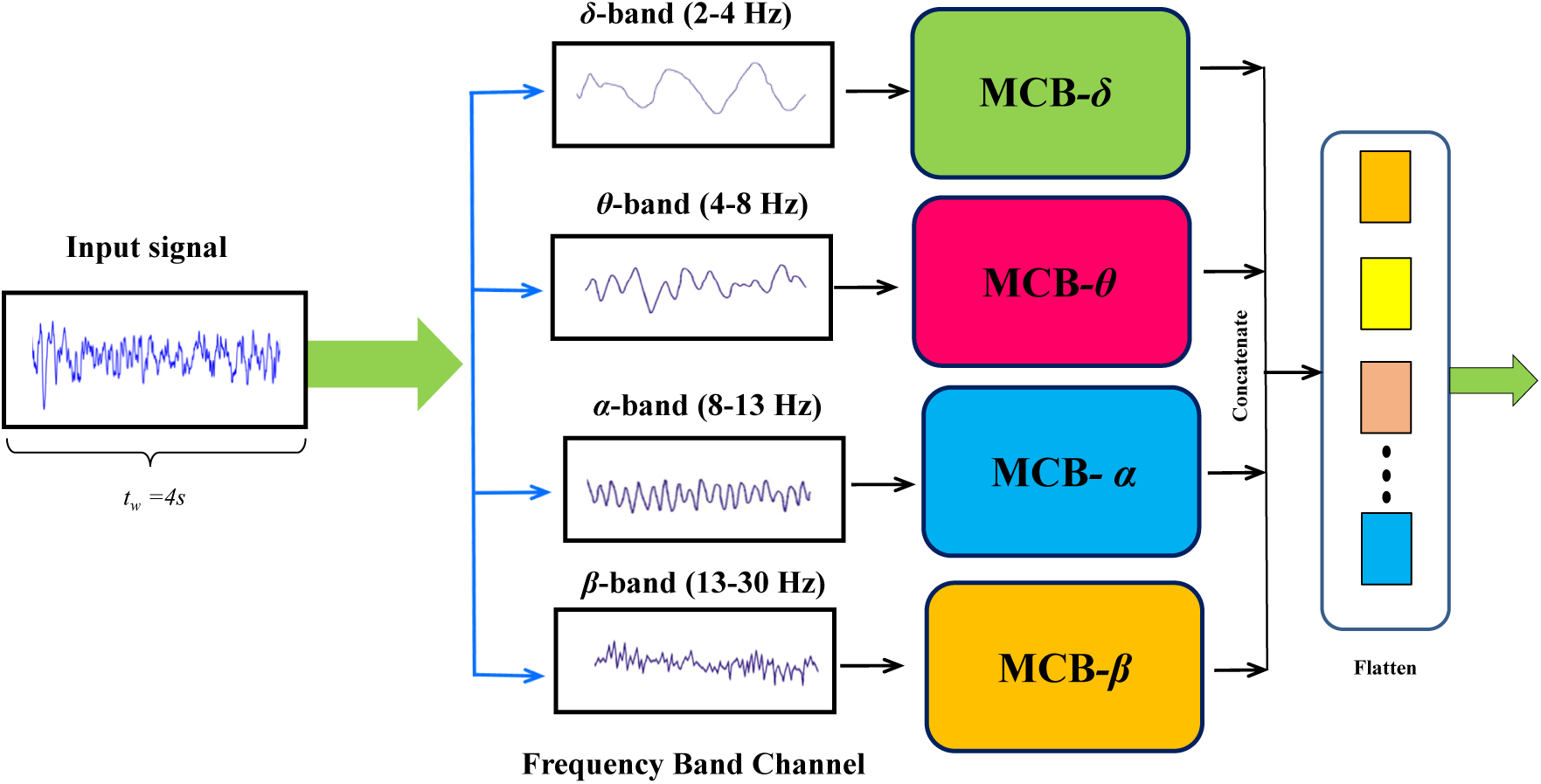
The architecture of multi-scale CNN (MSCNN) network architecture which consists of multi-scale convolution blocks (MCBs) as shown in Fig. 5.

Since our input EEG data representation is 1D time-series signal, 1D-CNN has been employed which is relatively easier to train and offers minimal computational complexity compare to its 2D counterparts whilst achieving state-of-the-art performance [57]. The convolution layer consists of 1D-convolution filters of a specified kernel stride which perform convolution operations sliding along the time axis of EEG signal to obtain feature maps and time-frequency information of the time series data [58, 59]. In 1D-CNN, forward propagation can be expressed as follows:

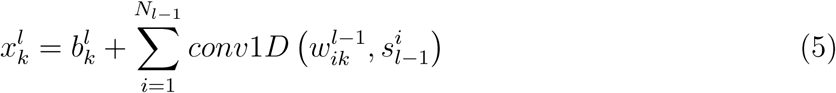

where 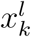 is the input; 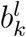 represents the bias of *k*^*th*^ feature information in *l*^*th*^ layer; 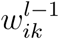 is defined as the connecting weight between *i*^*th*^ feature of the *l* − 1^*th*^ layer and *k*^*th*^ feature of the *l*^*th*^ layer; 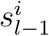 represents output of the *i*^*th*^ feature of *l* − 1^*th*^ layer; *conv*1*D* denotes convolution operation. The intermediate output 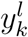 can be obtained passing 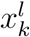 through the activation function *f* (•) as

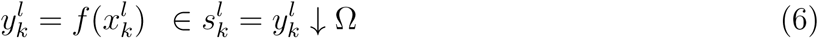

Where 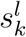 is the output of *k*^*th*^ feature information in *l*^*th*^ layer; Ω denotes the scalar factor of the down sampling operation ↓; rectified linear units (ReLU) *f* (*x*) = *max*(0, *x*) has been chosen as the activation function. During back-propagation, 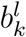 and 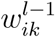 are updated when weight and bias sensitivities are determined by minimizing the error value *E*. The learning rate *η* can be defined as

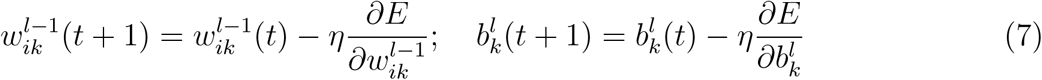

The convolutional layer is followed by the pooling layer or down-sampling layer to reduce the spatial size of the representation and the number of network parameters while preserving important and relevant features information. It aggregates neighboring values in the feature map by taking the average (average pooling) or maximum (maximum pooling) of the feature information. However, max pooling is more efficient in preserving the main features of the previous layer after down-sampling an input representation. Thus max pooling has been adopted in the present study.

### 4.2 Multi-scale convolution block (MCB)

During the convolution operation, different kernel size can extract different spatial feature map. For example, relatively large kernel size captures the overall feature. However, it may miss the relevant and important fine-grain feature information. In such case, a relatively small kernel size can effectively retain fine-grain information [60, 61]. Thus, in order to improve the performance of MI classifier, a multi-scale convolution consist of various kernel sizes can be an effective strategy to preserve both fine-grain high frequency localized information as well as low-frequency overall representations for the various frequency bands of EEG signal [31, 43].

Thus, a multi-scale convolution block (MCB) has been designed consists of three different kernel size in the convolution process. The network architecture of MCB has been shown in Fig. 5. In MCB, convolution block *λ*_*S*_ represents relatively small kernel size 1 × 1 which can effectively capture fine-grain localize information of the EEG signal. The convolution block *λ*_*M*_ with medium kernel size 1×3 can capture relatively coarse grain feature information. Whereas, *λ*_*L*_ represents large kernel size 1 × 5 which can collect the overall feature map efficiently. These three blocks are then followed by max-pooling layer to further reduce the network parameters. In addition, max-pooling and convolution layer of 1 × 3 has been utilized to preserve important features [31, 43] as shown in Fig. 5. In MSCNN, EEG signal has been divided into four different frequency bands channels and passed through corresponding MCB blocks (i.e., MCB_*i*_, *i* = *δ, θ, α*, and *β*) as shown in Fig. 4. Finally, the multi-scale feature information has been obtained by concatenation operation. The network parameters of MCB architecture has been detailed in Table. 1. The proposed MSCNN network can extract feature information from EEG signals on multiple scales which can significantly improves the classification accuracy of the MI-BCI classifier.

**Figure 5:**
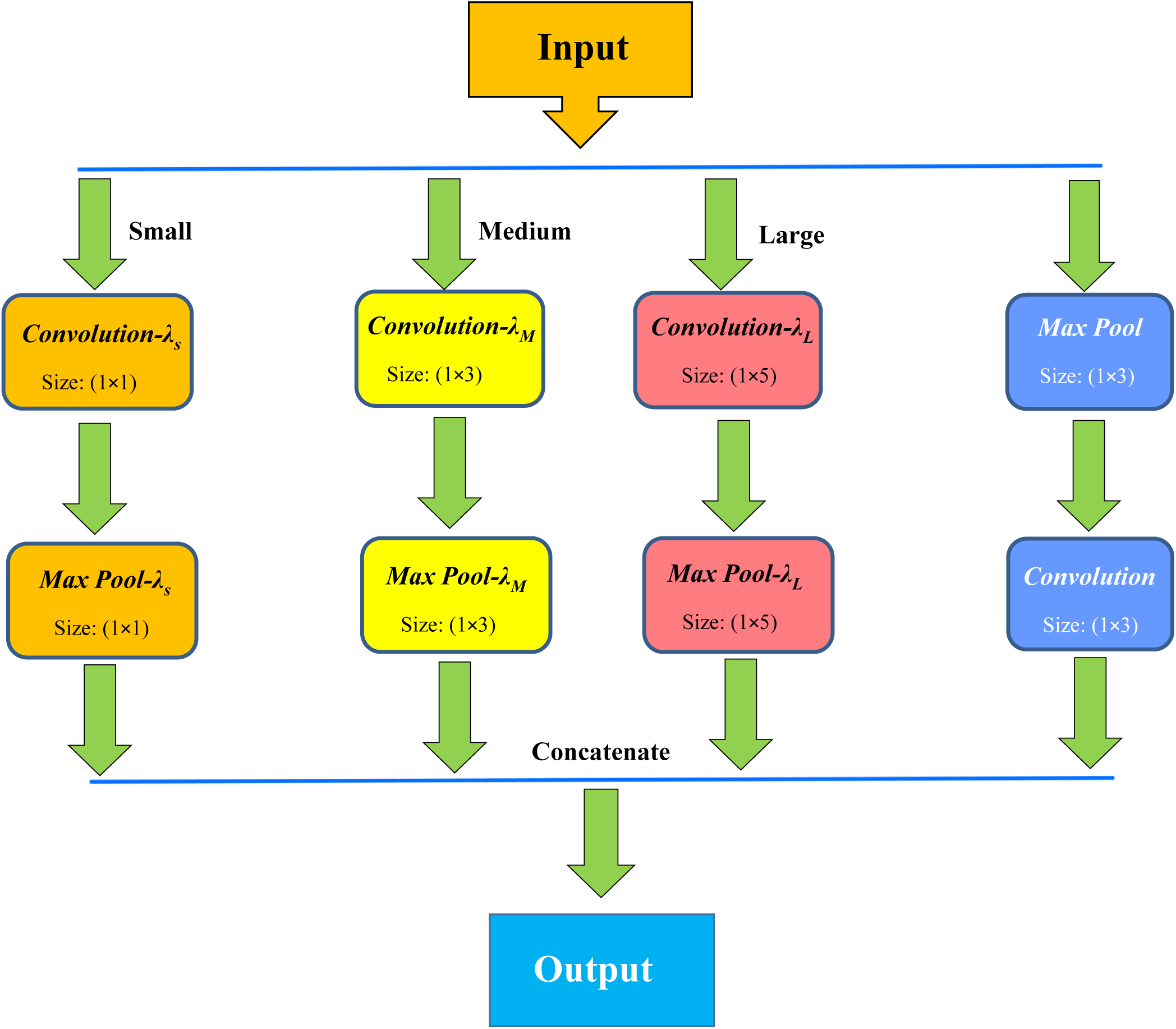
Network architecture of proposed MCB comprise of large (*λ*_*L*_), medium (*λ*_*M*_), and small (*λ*_*S*_) convolution scale for multi-scale feature extraction. See Table. 1 for network parameters of MCB.

**Figure 6:**
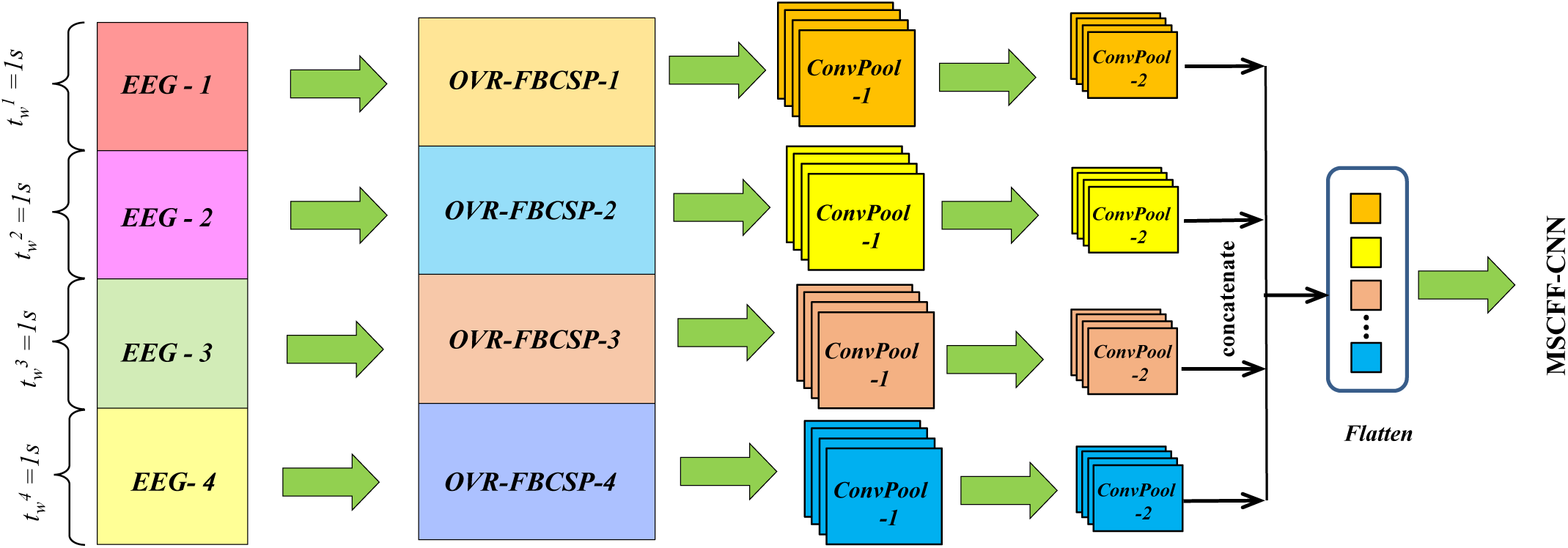
Schematic of network structure for proposed OVR-FBCSP CNN block. See Table. 2 for network parameters of OVR-FBCSP CNN.

**Table 1:** Network parameters of MCB architecture as shown in Fig. 5.

| Layer | Filter size | No. of Filters | Stride | Padding | Feature map size |
| --- | --- | --- | --- | --- | --- |
| Convolution Layer-$\lambda_S$ | $1 \times 1$ | 2 | 1 | SAME | $(1 \times 1000) \times 2$ |
| Max-Pooling Layer-$\lambda_S$ | $1 \times 1$ | - | 4 | SAME | $(1 \times 250) \times 2$ |
| Convolution Layer-$\lambda_M$ | $1 \times 3$ | 2 | 1 | SAME | $(1 \times 1000) \times 2$ |
| Max-Pooling Layer-$\lambda_M$ | $1 \times 3$ | - | 4 | SAME | $(1 \times 250) \times 2$ |
| Convolution Layer-$\lambda_M$ | $1 \times 5$ | 2 | 1 | SAME | $(1 \times 1000) \times 2$ |
| Max-Pooling Layer-$\lambda_M$ | $1 \times 5$ | - | 4 | SAME | $(1 \times 250) \times 2$ |
| Max-Pooling Layer | $1 \times 3$ | - | 4 | SAME | $(1 \times 250)$ |
| Convolution Layer | $1 \times 3$ | 2 | 1 | SAME | $(1 \times 250) \times 2$ |
| Flatten layer (after concat) | - | - | - | - | $1 \times 2000$ |

### 4.3 OVR-FBCSP CNN

The proposed model includes CNN layer after OVR-FBCSP output for each 1 *sec* time window called OVR-FBCSP CNN to extract spatial features as illustrated in Fig. 6. Each of the OVR-FBCSP CNN consists of two convolution-pooling layers. The feature output of size 12 × 12 for each time window from OVR-FBCSP has been passed through the 2D-convolution layer. The output of feature map 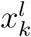 after 2D-convolution operation can be expressed as

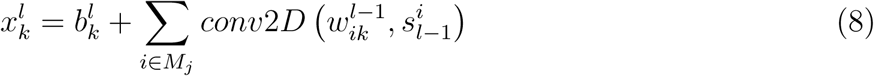

where *M*_*j*_ represents input feature collection; *conv*2*D* denotes 2D-convolution operation. The activation function ReLU has been used. The convolution layer is followed by max-pooling layer in order to reduce the size of the feature map. The stride of the pooling kernel has been chosen as 2. Additionally, the zero-padding method has been employed to preserve the edge information and size of the spatial feature map. Finally, the output of the max-pooling layer has been resized through flatten layer which produces 1 × 432 array. The main network parameters of OVR-FBCSP CNN architecture have been listed in Table. 2. The local features extracted by MSCNN and OVR-FBCSP have been concatenated together to form the global feature. It has been connected to convolution and then max-pooling layer as shown in Fig. 3.

**Table 2:**
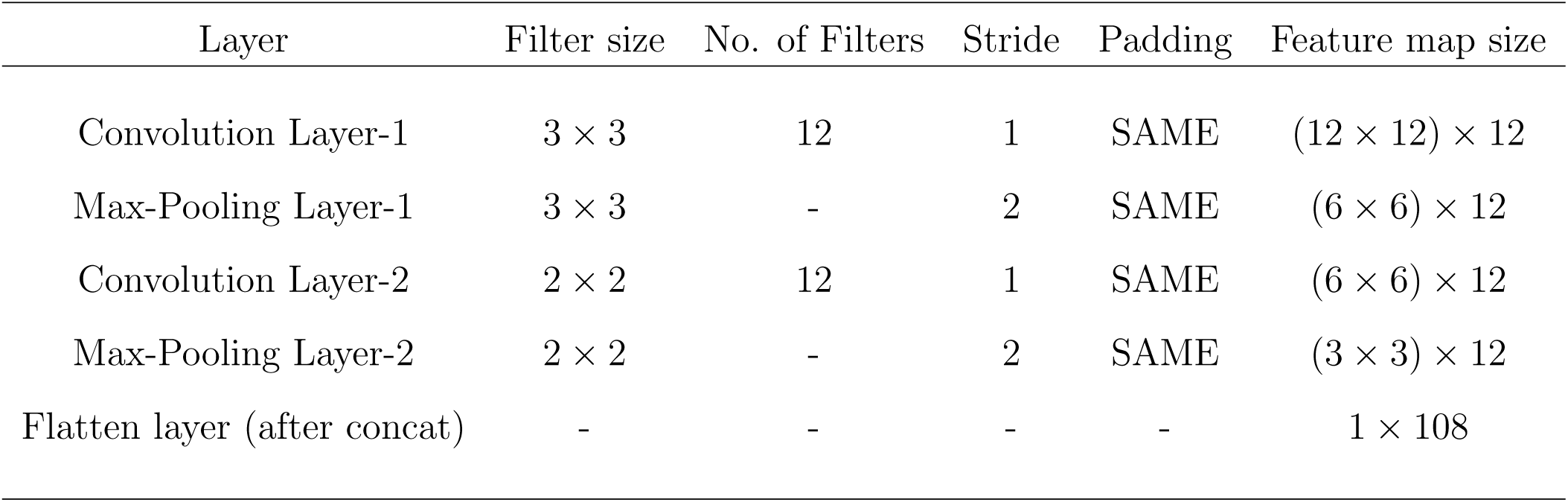
Network parameters of single branch OVR-FBCSP CNN architecture as shown in Fig. 6.

**Table 3:** Network parameters of proposed MSFFCNN model as shown in Fig. 3.

| Block/Layer | Kernel size | No. of Kernal | Stride | Padding | Feature map size |
| --- | --- | --- | --- | --- | --- |
| MSCNN | - | - | - | - | $4 \times (1 \times 2000)$ |
| OVR-FBCSP CNN | - | - | - | - | $4 \times (1 \times 108)$ |
| Convolution Layer | $1 \times 3$ | 2 | 1 | SAME | $(1 \times 8432) \times 2$ |
| Max-Pooling Layer | $1 \times 3$ | - | 4 | SAME | $(1 \times 2108) \times 2$ |
| FC-1 | - | - | - | - | 1024 |
| Dropout layer | - | - | - | - | - |
| FC-2 | - | - | - | - | 512 |
| FC-3 | - | - | - | - | 256 |
| Output layer | - | - | - | - | 4 |

### 4.4 FC layers

The extracted spatial features obtained from the max-pooling layer are passed through FC layers consisting of by 3-layer feedforward neural network as shown in Fig. 3. Finally, it is connected to the Softmax layer containing four neurons to predict the final multi-classification result. The proposed MSFFCNN model has been fine-tuned to increase accuracy of the MI-BCI classifier by continuous experimental adjustment. The overall network parameters of proposed MSFFCNN model has been outlined in Table. 3.

### 4.5 Training optimization

The fully connected layer utilizes ReLU as the activation function in hidden layers which helps to accelerates the optimization process of the network. It provides better classification accuracy compared to other activation functions for MI-BCI application [31]. The softmax function has been utilized to obtain exponential probability distribution of 4 different MI-BCI classification tasks in output layer which can be expressed as:

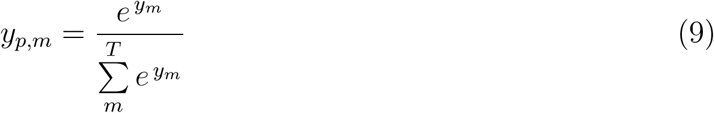

where *T* represents the total number of classes; *m* represents the index of corresponding classes. Additionally, cross-entropy loss function has been utilized during training to optimize the model. The cross-entropy *L*_*CE*_ can be expressed as: 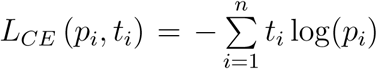 where *n* is total number of classes; *t*_*i*_ is the truth label; *p*_*i*_ is the Softmax probability for *i*^*th*^ class. In addition, Adam [62] optimization scheme has been implemented to minimize the difference between probabilistic cross-entropy loss. Moreover, the dropout technique [63] has been employed to prevent over-fitting and accelerate the training procedure.

## 5. Transfer learning modeling

Generally, CNN based MI-BCI classification algorithms contain large numbers of trainable model parameters which requires significant amount of training data and leads to an increase in computation time [36, 37]. In order to overcome the aforementioned issues, transfer learning can be an efficient strategy utilizing pre-trained weights from different subject cases. [36, 37, 39, 40, 41] However, transfer learning can be challenging in the MI-BCI system due to substantial inter-subject variability between different subjects [44, 40]. Therefore, adaptation schemes require special attention for fine-tuning the model parameter prudently to establish an efficient transfer learning-based MI-BCI classifier [39, 40, 41]. In the present study, three different classification procedure including subject-specific, subject-independent and subject-adaptive classification method has been employed which have been detailed in subsequent sections.

### 5.1 Subject-specific classification model (MSFFCNN-1)

In the present study, at first, the conventional subject-specific classification approach has been employed where the proposed model is trained on the target subject and the performance of the model has been evaluated on the same subject. The trained model based on subject-specific classification is the first baseline model for the present study and it has been named MSFFCNN-1.

### 5.2 Subject-independent classification model (MSFFCNN-2)

For the second approach, Subject-independent classification has been adopted where the model has been trained with all possible available training data except the target subject. During validation, LOSO cross-validation [37] procedure has been utilized. The subject-independent classification-based model has been considered as the second baseline model. In the present work, it has been referred to as MSFFCNN-2.

### 5.3 Subject-adaptive classification model (MSFFCNN-TL)

Additionally, to further improve the performance of the subject-independent classification model, subject-adaptive classification approach has been employed where the model has been fine-tuned for a particular subject in a pre-trained model (MSFFCNN-2) with a different fraction of target subject data (see section 5.4 for datasets division) to study the influence of various degrees of adaptation *ξ* on the accuracy and performance of the classifier. In this study, the subject-adaptive classification-based model has been termed as MSFFCNN-TL. The proposed MSFFCNN-TL has two variants MSFFCNN-TL-1 and MSFFCNN-TL-2 based on two different adaptation configurations *AC* − 1 and *AC* − 2, respectively as shown in Fig. 7. In the first adaptation configuration *AC* − 1, the fully connected (FC) layers have been adopted and optimized, whereas, rest of the network parameters have been kept unchanged. In the second adaptation configuration *AC* − 2, the last conv-pool block and fully connected layer have been retrained using the adaptation data while the rest of the network parameters have been kept unchanged. These two proposed adaptation configurations have been illustrated in Fig. 7-(a). The degree of adaptation *ξ* can be defined as the fraction of training data to fine-tune the model for each subject-adaptive configuration as shown in Fig. 7 -(b). The numbers of required trainable network parameters Ψ_*AS*_ for two adaption configurations have been outlined in Fig. 7 -(c). Additionally, learning rate *η* has been scaled down to avoid clobbering the initialization [64]. From the Eq. 7, one can assume 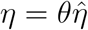 where 0 *< θ* ≤ 1 is the scaling factor of *η*:

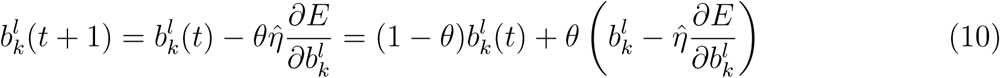

For *θ <* 1, the model accepts scaled-down weighted adaptation. In the present study, different scaling factors *θ* has been considered to obtain the optimal choice of the learning rate for efficient adaptation and enhance the performance of the classifier (see section 6.3.1).

**Figure 7:**
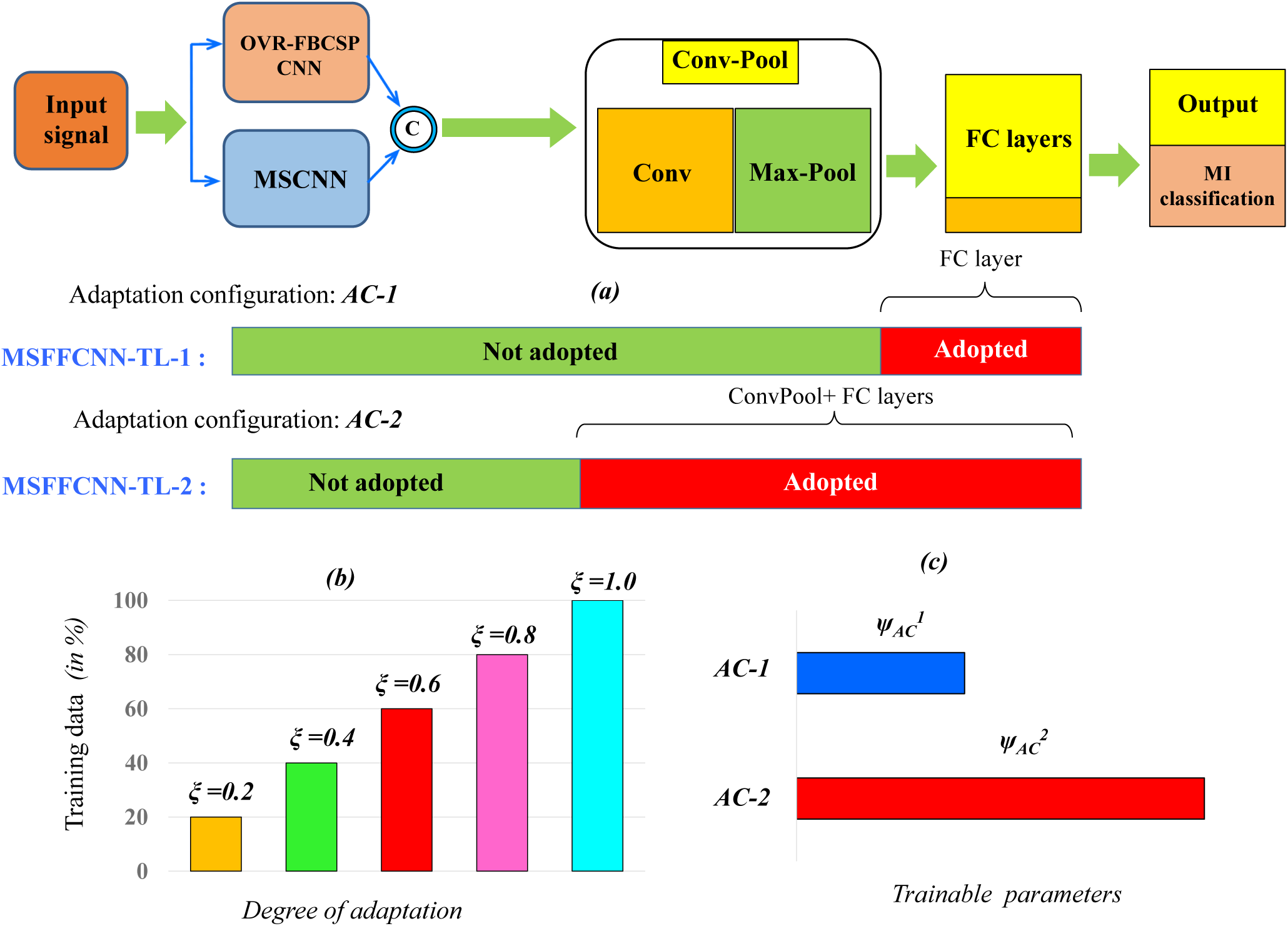
Illustrations of (a) two different adaption configurations *AC* − 1 and *AC* − 2 for the proposed subject adaptive classification models MSFFCNN-TL-1 and MSFFCNN-TL-2, respectively; (b) different degree of adaption *ξ*; (c) numbers of required trainable network parameters Ψ_*AS*_ for adaption configurations.

**Figure 8:**
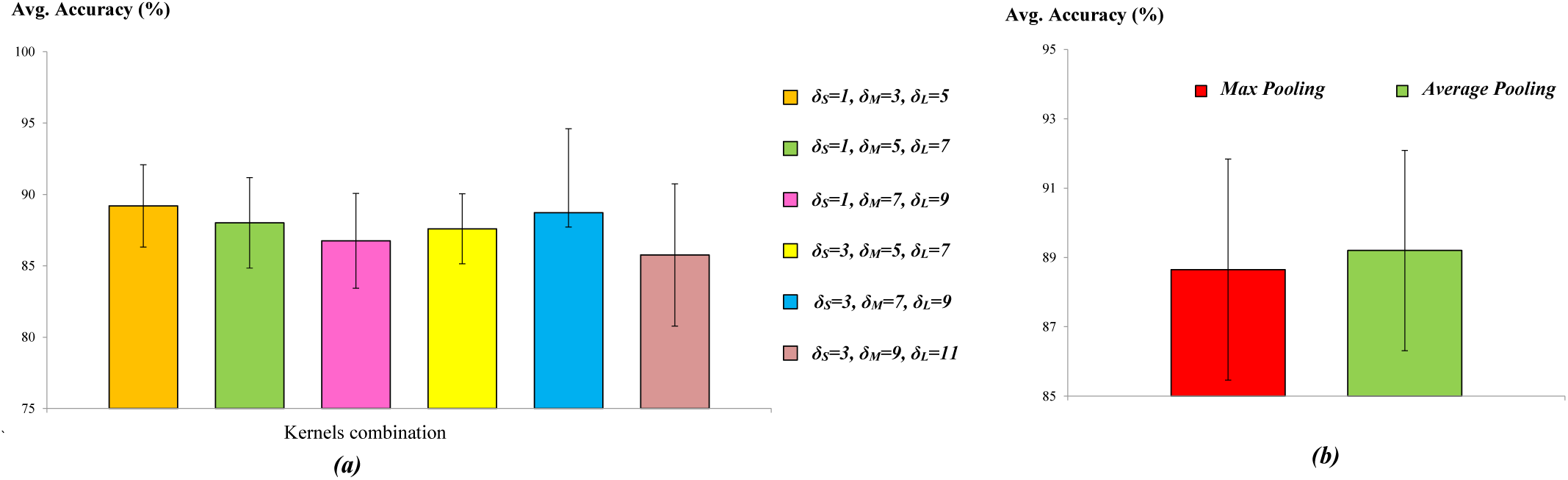
Comparison of average accuracy with SE (in %) from MSFFCNN-1 for (a)various combinations of kernel sizes; (b) average pooling and max pooling in HCB.

### 5.4 Dataset arrangement for transfer learning

In this section, the subdivision of EEG data during training and evaluation of the transfer learning models has been detailed. From the BCI competition IV-2a EEG dataset, the first session (identifier: *T*) is used to train the classifier. The second session (identifier: *E*) has been strictly utilized for evaluating the corresponding trained classifier. For subject-specific classification model MSFFCNN-1, first and second sessions have been considered for training and evaluation, respectively for a particular subject. In subject-independent classification, first session datasets from all subjects except the target subject have been utilized to train the MSFFCNN-2 model. For subject-adaptive classification models MSFFCNN-TL, the model corresponding to minimum validation loss in the subject independent classification across all nine subjects has been chosen as the pre-trained model. To fine-tune the pre-trained MSFFCNN-2 model, the different fractions of training data ranging from 20% to 100% in steps of 20% have been considered. For a particular subject, the subject-adaptive classification model has been validated with the corresponding evaluation test dataset.

## 6. Results and discussions

In this section, the accuracy and performance of the subject-specific, subject-independent, and subject-adaptive classification models have been discussed and compared with several existing state-of-the-art methods. The performance of the proposed model has been evaluated by classification accuracy 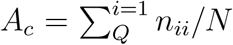 which can be obtained from the confusion matrix by the ratio of the sum of diagonal elements 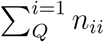 to the total number of samples *N*. For a fair comparison, a similar training strategy has been applied for proposed MSFFCNN-1, MSFFCNN-2, and MSFFCNN-TL models. During training the models, batch size of 24 has been considered which attains the highest classification accuracy and optimizes the convergence speed. The network has been trained for 350 epochs. In the second FC layer, a dropout probability value of 0.5 has been prescribed. The models have been implemented in Keras API with TensorFlow as backend. The model has been trained and tested using an Intel Core i7 CPU with a single NVIDIA GeForce RTX 2080 8 GB GPU. The average *A*_*c*_ value and standard error 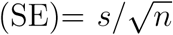 (*s* is the sample standard deviation; *n* is the sample size) are calculated across all subjects.

### 6.1 Result for subject-specific classification

In this section, various network optimizations and corresponding average classification accuracy have been reported for subject-specific classification MSFFCNN-1 model.

#### 6.1.1. Effect of convolution kernel size

As mentioned earlier, different kernel sizes of the convolution layer can capture distinct features of the EEG signal. Thus, various combinations of convolution blocks (i.e, *λ*_*S*_, *λ*_*M*_, and *λ*_*L*_ in Fig. 5) in MCB are considered to study the effect of kernel sizes on the performance and accuracy of the proposed model as reported in Table 4. With increasing kernel size, in particular, for combinations (*δ*_*S*_ = 3, *δ*_*M*_ = 7, *δ*_*L*_ = 9) and (*δ*_*S*_ = 3, *δ*_*M*_ = 9, *δ*_*L*_ = 11) classification accuracy have been achieved to 88.72% and 85.76%, respectively as shown in Fig. 8 -(a). For combinations (*δ*_*S*_ = 1, *δ*_*M*_ = 7, *δ*_*L*_ = 9) and (*δ*_*S*_ = 3, *δ*_*M*_ = 5, *δ*_*L*_ = 7) classification accuracy drops down to 86.75% and 87.59%, respectively. For the combination of relatively small kernel sizes, classification accuracy improves.

**Table 4:** Average classification accuracy with SE (in %) from MSFFCNN-1 model for various combinations of kernel sizes in MCB.

| Kernel size |  |  | Avg. Accuracy | SE |
| --- | --- | --- | --- | --- |
| Small ($\delta_S$) | Medium ($\delta_M$) | Large ($\delta_L$) | | |
| 1 | 3 | 5 | **89.20** | $\pm 2.89$ |
| 1 | 5 | 7 | 88.01 | $\pm 3.17$ |
| 1 | 7 | 9 | 86.75 | $\pm 3.33$ |
| 3 | 5 | 7 | 87.59 | $\pm 2.45$ |
| 3 | 7 | 9 | 88.72 | $\pm 5.89$ |
| 3 | 9 | 11 | 85.76 | $\pm 4.98$ |

**Table 5:** Average classification accuracy with SE (in %) obtained from MSFFCNN-1 for average pooling and max pooling

| **Average Accuracy with SE (%)** |  |
| --- | --- |
| Average Pooling | Max pooling |
| 88.65 ±3.19 | 89.20 ±2.89 |

After considering all combinations, the best classification result has been obtained corresponds to *δ*_*S*_ = 1, *δ*_*M*_ = 3, *δ*_*L*_ = 5 with an average classification accuracy of 89.20%. Thus, in the present study, combinations of Ω_*L*_ = 5, Ω_*M*_ = 3, and Ω_*S*_ = 1 has been selected. Such combination is found to be computationally most efficient with with 1.19% accuracy gain compared to Ω_*S*_ = 1, Ω_*M*_ = 5, and Ω_*L*_ = 7 (see Table 9). Moreover, such combination achieves SE value of 2.89% which is 0.28% lower than Ω_*S*_ = 1, Ω_*M*_ = 5, and Ω_*L*_ = 7. Thus, *δ*_*S*_ = 1, *δ*_*M*_ = 3, *δ*_*L*_ = 5 in MCB provides the best performance in-terms of both classification accuracy and speed from MSFFCNN-1 model.

#### 6.1.2 Influence of pooling type

From the previous section, it has been concluded that the combination of *δ*_*S*_ = 1, *δ*_*M*_ = 3, *δ*_*L*_ = 5 provides the best accuracy. Thus, corresponding pooling window size of 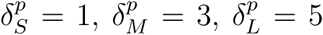 has been prescribed in MCB. Two different types of pooling including max pooling and average pooling have been considered to explore the effect of pooling type on the average classification of MSFFCNN-1 as listed in Table. 5. From the comparison, one can see that max-pooling provides the best classification accuracy with 0.55% improvement over average-pooling as depicted in Fig. 8 -(b). Moreover, max-pooling provides the smallest SE value of 2.89% that is 0.30% lower than average pooling. Thus, max-pooling has been adopted in all four models which provides the better capability of highlighting features conducive to the MI-BCI classification task.

### 6.2 Result for subject-independent classification

In this section, the classification accuracy for each subject and average classification accuracy across all 9 subjects has been reported for subject-independent classification model MSFFCNN-2 compared with MSFFCNN-1 and other baseline models.

#### 6.2.1 Comparison with baseline models

In order to evaluate the performances of the proposed MSFFCNN variants, classification accuracy is compared with commonly used traditional ML models as MI-BCI classifier including SVM and LDA as the baseline models. In addition, a standard CNN have been used as a baseline DL model. The standard CNN network structure consists of five conv-pool layers which are relatively deeper than proposed MSFFCNN models. For fair comparison, model hyper-parameters were kept as consistent as possible with the proposed models. As outlined in Table 6, both MSFFCNN-1 and MSFFCNN-2 have achieved better accuracy for all subjects with an average accuracy of 89.20% ± 2.89% and 91.61% ± 2.49% compared to baseline models. Compared to LDA and SVM, MSFFCNN-1 has gained 20.43% and 11.05% improvement in average *A*_*c*_, respectively. As shown in Fig. 9 -(b), both MSFFCNN-1 and MSFFCNN-2 have demonstrated superior performance with accuracy improvement of 13.65% and 14.15% over the standard CNN model. Additionally, the MSFFCNN models have relatively low SE value indicating better capability and generalization in representing EEG data from different subjects compared to other baseline models. The comparison illustrates the advantages of using the DL model for strong feature extraction capability among different subjects in EEG-based MI-BCI system compared to traditional ML methods.

**Table 6:**
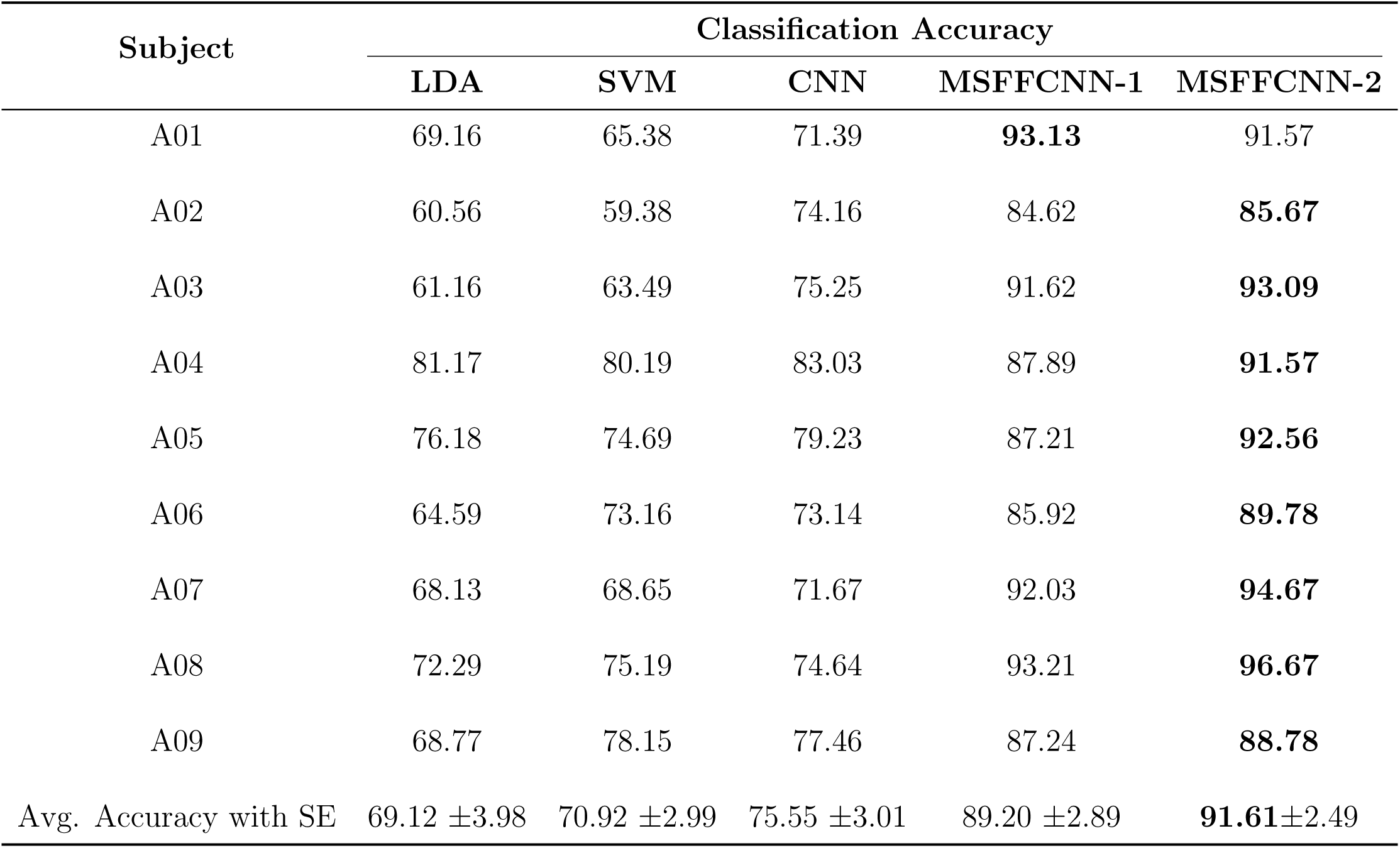
Classification accuracy for each subject and average accuracy with SE from LDA, SVM, standard CNN, subject-specific model MSFFCNN-1, and subject-independent model MSFFCNN-2. The bold indicates the best result.

**Figure 9:**
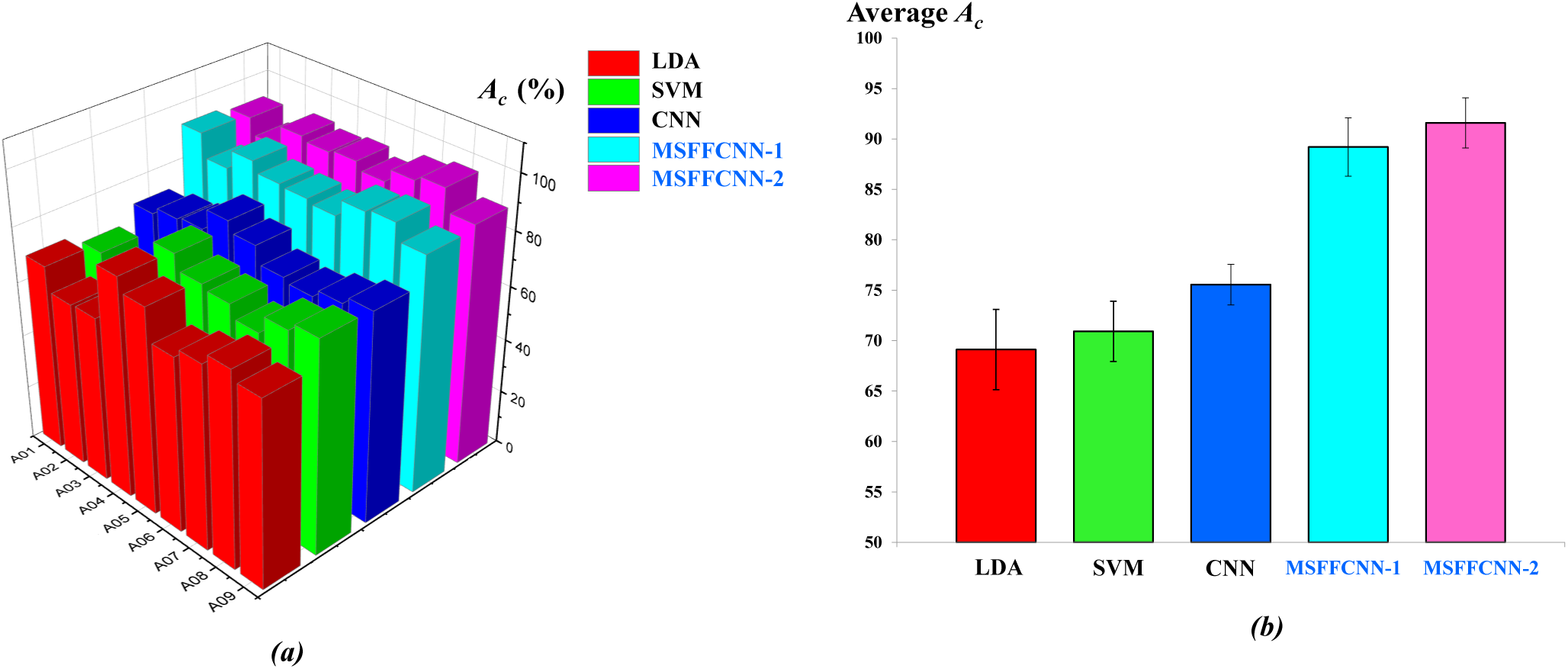
Comparison bar chart of (a) *A*_*c*_ for each subject; (b) average *A*_*c*_ with SE (in %) between LDA, SVM, CNN, and proposed MSFFCNN models.

#### 6.2.2 Comparison of subject-dependent and subject-independent models

Comparing the *A*_*c*_ values, MSFFCNN-2 has achieved better average *A*_*c*_ with 2.41% improvement over MSFFCNN-1 as illustrated in Fig. 9-(b). For subject A01, MSFFCNN-1 has demonstrated better performance with *A*_*c*_ of 93.913% compared to 91.57% achieved by MSFFCNN-2. However, MSFFCNN-2 has demonstrated better accuracy for all other subject classes, in particular there are 5.35%, 3.86%, and 3.46% improvement in A05, A06, and A08, respectively compared to MSFFCNN-1. The comparison indicates that relatively large training data can significantly increase the performance of MSFFCNN-2. Furthermore, MSFFCNN-2 has achieved a lower SE value of 2.49% indicating better representation and generalization of the classifier than MSFFCNN-1.

### 6.3 Result for subject-adaptive classification

This section reports the performance of subject-adaptive classification model MSFFCNN-TL for two different adaptation configurations *AC* − 1, *AC* − 2. Various parametric study considering different learning rate scale factor *θ* and degree of adaptation *ξ* on the accuracy of MSFFCNN-TL classifier has been explored in the following subsections.

#### 6.3.1 Influence of learning rate scale factor *θ*

As mentioned earlier, different *θ* can have significant impact on the performance of the MSFFCNN-TL model, thus in the current work six different *θ* = 1, 0.5, 0.1, 0.05, 0.025, and 0.01 have been studied considering five different degree of adaption *ξ* = 0.2, 0.4, 0.6, 0.8, and 1.0 for each *θ* as outlined in Tables 7-8 where bold highlights the best adaptation configuration for a particular *θ*. For *θ* = 1, the learning rate of the MSFFCNN-TL model is the same as the other two models. In such a case, the best average *A*_*c*_ has been obtained as 91.79% for the full degree of adopted *AC* − 2 configuration as shown in Table 7. However, partial degree of adaptation *ξ* = 0.8 in *AC* − 2 provides the best *A*_*c*_ = 92.36% for *θ* = 0.5. With further reduction of *θ* = 0.1 there is 0.88% improvement in accuracy compared to *θ* = 0.5 for *ξ* = 1.0. Similarly, for *θ* = 0.05 and 0.025, accuracy has been improved to 93.69% and 93.84% for full adaptaion *ξ* = 1.0, respectively as shown in Table 8. It is noteworthy to mention, the best accuracy results for *θ* = 0.05 and 0.025 have been achieved for *AC* − 2. However, the overall best average *A*_*c*_ = 94.06% has been obtained for the lowest *θ* = 0.01 with 2.27% increase compared to *θ* = 1.0 indicating importance of tuning down *θ* to improve the performance of the classifier significantly.

**Table 7:**
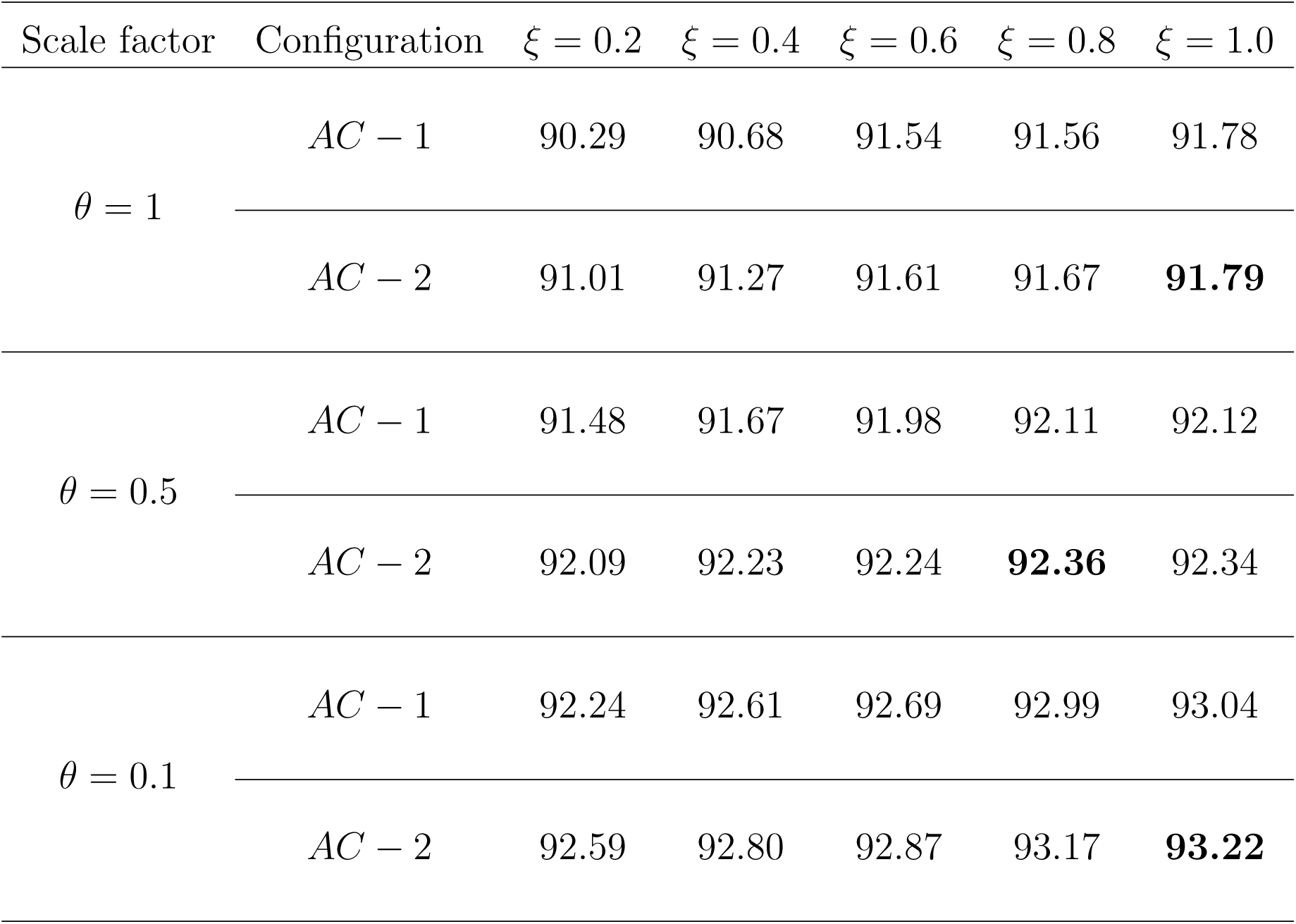
Average *A*_*c*_ obtained from MSFFCNN-TL model for *θ* = 1, 0.5, and 0.1 considering different *ξ* for *AC* − 1 and *AC* − 2 configurations. The bold highlights the best adaptation configuration for a particular *θ*.

**Table 8:** Average *A*_*c*_ obtained from MSFFCNN-TL model for *θ* = 0.05, 0.025, and 0.01 considering different *ξ* for *AC* − 1 and *AC* − 2 configurations. The bold highlights the best adaptation configuration for a particular *θ*.

| Scale factor | Configuration | $\xi = 0.2$ | $\xi = 0.4$ | $\xi = 0.6$ | $\xi = 0.8$ | $\xi = 1.0$ |
| --- | --- | --- | --- | --- | --- | --- |
| $\theta = 0.05$ | $AC - 1$ | 93.17 | 93.21 | 93.38 | 93.57 | **93.69** |
| | $AC - 2$ | 92.98 | 93.08 | 93.45 | 93.52 | 93.67 |
| $\theta = 0.025$ | $AC - 1$ | 93.21 | 93.39 | 93.67 | 93.72 | **93.84** |
| | $AC - 2$ | 93.01 | 93.07 | 93.24 | 93.34 | 93.69 |
| $\theta = 0.01$ | $AC - 1$ | 93.17 | 93.23 | 93.35 | 93.69 | 93.72 |
| | $AC - 2$ | 93.18 | 93.36 | 93.98 | 94.01 | **94.06** |

**Table 9:** Comparison of average *A*_*c*_, *E*_*f*_, and *t*_*c*_ between MSFFCNN-1, MSFFCNN-2, MSFFCNN-TL-1 (*AC* − 1), and MSFFCNN-TL-2 (*AC* − 2) for some specific *θ* and *ξ*.

| Model | Configuration | $\theta$ | $\xi$ | Avg $A_c$ | $E_f$ | $t_c$ (in sec) |
| --- | --- | --- | --- | --- | --- | --- |
| MSFFCNN-1 | - | 1 | - | 89.20 | 4.34 | 9.78 |
| MSFFCNN-2 | - | 1 | - | 91.61 | 3.87 | 138 |
| MSFFCNN-TL-1 | $AC - 1$ | 0.5 | 0.6 | 91.98 | 3.43 | 7.83 |
| MSFFCNN-TL-2 | $AC - 2$ | 0.1 | 0.4 | 92.80 | 3.01 | 6.76 |
| MSFFCNN-TL-1 | $AC - 1$ | 0.025 | 0.8 | 93.72 | 2.67 | 8.78 |
| MSFFCNN-TL-2 | $AC - 2$ | 0.01 | 1 | 94.06 | 1.43 | 9.55 |

In Fig. 10, average *A*_*c*_ has been plotted as function of *ξ* for different *θ*. For *θ* ≤ 0.1, *A*_*c*_ increases with decreasing *θ* for particular *ξ* as shown in Fig. 10-(a) with *AC* − 2 provides best result. The result indicates that *AC* − 2 is suitable for better adaption for relatively high *θ*. However, for *θ >* 0.1, *AC* − 1 can be efficient, in particular, 0.6 ⩽*ξ* ⩽ 1.0 as illustrated in Fig. 10-(b) indicating better adaption for lower learning rate even for fewer number of adaptation network parameters. Overall, with higher degree of adaption, both configurations tend to perform better.

**Figure 10:**
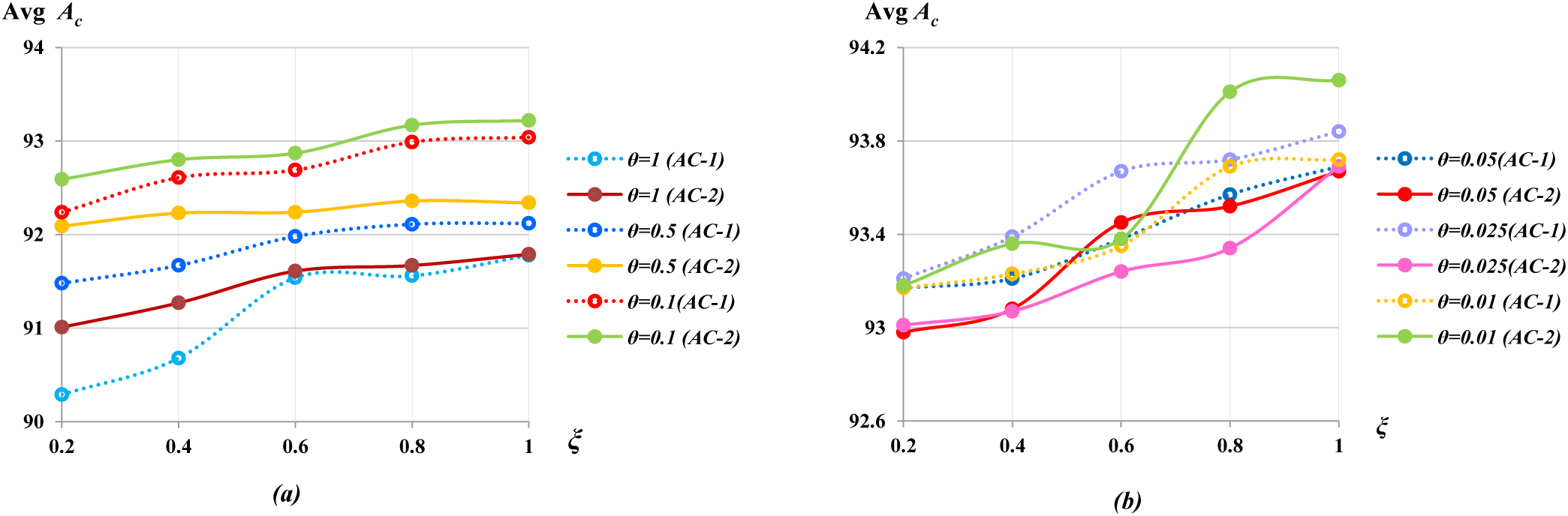
Influence of *θ* on the average *A*_*c*_ (in %) of the MSFFCNN-TL model for (a) *θ* = 1, 0.5, 0.1 ; (b) *θ* = 0.05, 0.025, 0.01 in the range 0.2 ⩽*ξ* ⩽ 1.0 considering both *AC* − 1 and *AC* − 2 configurations.

**Figure 11:**
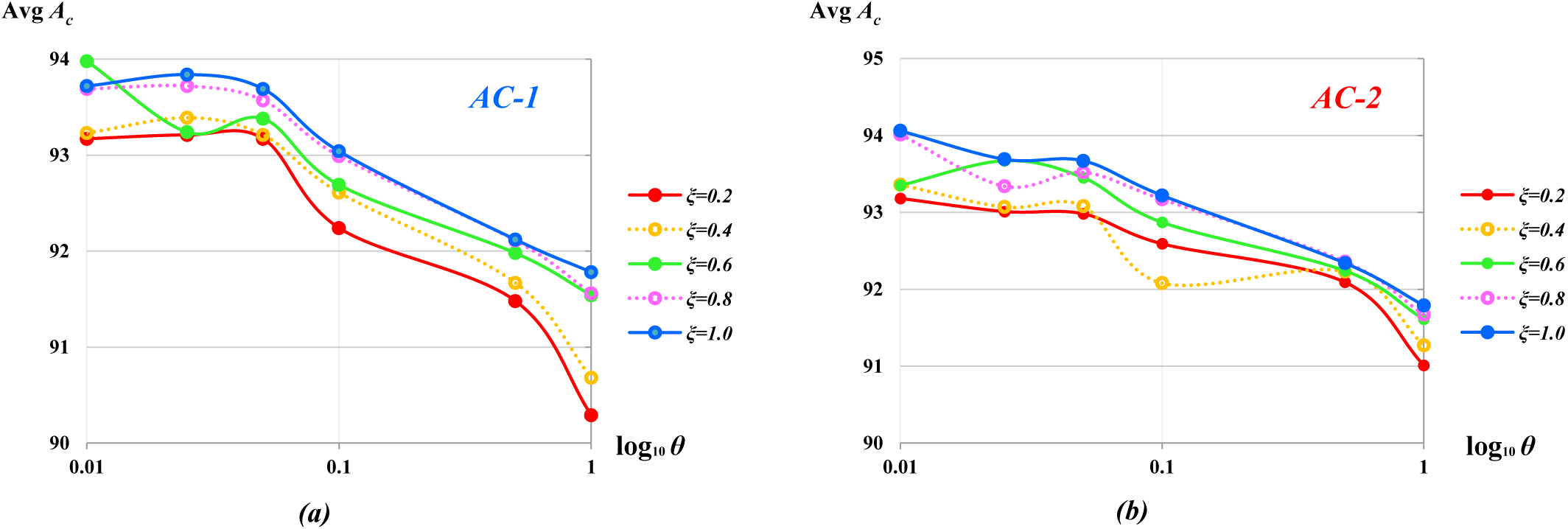
Influence of degree of adaption *ξ* on the classification accuracy *A*_*c*_ (in %) of the MSFFCNN-TL model for (a) *AC* − 1; (b) *AC* − 2 configurations in the range 0.01 ⩽ *θ* ⩽ 1.0.

#### 6.3.2 Influence of degree of adaptation ξ

In this section, the influence of *ξ* on *AC* of the MSFFCNN-TL classifier has been explored in the range 0.01 ⩽*θ* ⩽ 1.0 as shown in Fig.11. For *AC* − 1 configuration, increasing *ξ* for a particular *θ* increases average *A*_*c*_ at least in the range 0.025 ⩽ *θ* ⩽ 1.0 as depicted in Fig. 10-(a). For *ξ* = 0.6, the accuracy the classifier improves for *θ <* 0.05. Whereas, for *AC* − 2, the effect of relatively low *ξ* = 0.4 on the performance of the classifier is more prominent in relatively high *θ >* 0.1 as shown in Fig. 10-(b). This is because the *AC* − 2 configuration has a higher trainable parameter than *AC* − 1 which may lead to over-fitting to some degree for higher *ξ*. The present study reveals that there exist two different regimes of *θ* for two different adaptation configurations for optimum performance of the MSFFCNN-TL classifier.

#### 6.3.4 Overall performance comparison of four models

This section compares the overall performance of MSFFCNN-1, MSFFCNN-2, MSFFCNN-TL-1 (*AC* − 1), and MSFFCNN-TL-2 (*AC* − 2) by evaluating average *A*_*c*_, final loss *E*_*f*_, and computation time for training *t*_*c*_ as listed in Table 9. The comparison demonstrates the superiority of subject-adaptive model by achieving performs better than subject-dependent and subject-independent counterpart even for *ξ <* 1 implying effectiveness of the proposed MSFFCNN-TL model by using less training data. MSFFCNN-TL-2 shows better performance than MSFFCNN-TL-1 for larger number of training sample, in particular, *ξ >* 0.6 since MSFFCNN-TL-2 has larger number of network parameters to adopt. Overall, MSFFCNN-TL-2 has achieved the highest average *A*_*c*_ = 94.06% which is a 4.86% and 2.45% improvement over MSFFCNN-1 and MSFFCNN-2, respectively. Comparing *E*_*f*_, MSFFCNN-1 reaches the maximum value of 4.34, whilst, MSFFCNN-TL for *θ* = 0.01 attains the lowest value of 1.43 indicating better learning capability between all four model variants. Additionally, comparison of *t*_*c*_ reveals that MSFFCNN-1 and MSFFCNN-TL variants take similar time-frame (within 10 *sec*) to train where MSFFCNN-1 takes slightly higher training time. With increase of *ξ*, training time increases. For same *ξ* and *θ*, configuration *AC* − 1 is faster compared to *AC* − 2 due to lower number of trainable parameters. Whereas, MSFFCNN-2 take significantly longer time to train with average *t*_*c*_ of 138 *sec* (or 2.3 *min*). The comparison demonstrates that the subject-adaptive model has better accuracy, learning capability, and adaptability for different subject classes with significantly less training data and training time.

### 6.4 Comparison Cohen’s kappa-coefficient

To evaluate the performance of proposed MI-BCI models, average Cohen’s kappa-coefficient (*κ*) has been calculated as [65]:

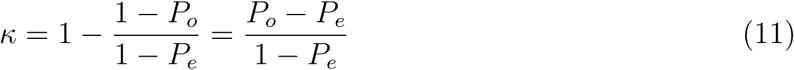

where *P*_*e*_ can be measured by summing the product of predicted numbers for each category and ground truth for each category divided by the square of the total number of samples; *P*_*o*_ denotes the total classification accuracy. For multi-class classification problem, *κ* value reflects the generalization of the model and consistency of classifying different subjects. In Table 10, *κ* values obtained from the MSFFCNN-2 and MSFFCNN-TL models have been compared with the state-of-art ML models such as FBCSP [66], channel-wise CNN (CW-CNN) [42], 3D CNN [67], channel-wise convolution with channel mixing (C2CM) [42], sequential updating semi-supervised spectral regression kernel discriminant analysis (SUSS-SRKDA) [68], deep restricted Boltzmann machine network with t-distributed stochastic neighbor embedding (RBMt-SNE) [69], stacked regularized linear discriminant analysis (SRLDA) [70], hybrid deep neural network (HDNN) [41], CNN-LSTM [34], EEG topographical representative CNN (ETRCNN) [71], and HDNN with transfer learning (HDNN-TL) [41] for BCI Competition IV 2a datasets. From the overall comparison, it can be seen that the proposed models outshine other ML models significantly in terms of *κ* values as shown in Fig. 12. Comparing average *κ*, MSFFCNN-TL has achieved the highest *κ* value of 0.88 which is 10.00% and 8.00% improvement over the state-of-the-art HDNN and ETRCNN models, respectively.

**Table 10:** Comparison of *κ* with several state-of-the-art ML models across all subjects. The bold denotes the best result from the corresponding model.

| Method | Subject |  |  |  |  |  |  |  |  |  |
| --- | --- | --- | --- | --- | --- | --- | --- | --- | --- | --- |
|  | A01 | A02 | A03 | A04 | A05 | A06 | A07 | A08 | A09 | Avg |
| FBCSP | 0.67 | 0.41 | 0.74 | 0.48 | 0.39 | 0.27 | 0.77 | 0.75 | 0.60 | 0.56 |
| CW-CNN | 0.81 | 0.47 | 0.82 | 0.56 | 0.50 | 0.26 | 0.87 | 0.75 | 0.69 | 0.64 |
| 3D CNN | 0.69 | 0.45 | 0.78 | 0.59 | 0.64 | 0.53 | 0.65 | 0.70 | 0.71 | 0.64 |
| C2CM | 0.83 | 0.53 | 0.87 | 0.55 | 0.50 | 0.27 | 0.86 | 0.77 | 0.72 | 0.65 |
| SUSS-SRKDA | 0.83 | 0.51 | 0.88 | 0.68 | 0.56 | 0.35 | 0.90 | 0.84 | 0.75 | 0.70 |
| RBN $t$-SNE | 0.82 | 0.48 | 0.76 | 0.66 | 0.50 | 0.53 | 0.78 | 0.86 | 0.89 | 0.70 |
| SRLDA | 0.84 | 0.55 | 0.90 | 0.71 | 0.66 | 0.44 | 0.94 | 0.85 | 0.76 | 0.74 |
| HDNN | 0.82 | 0.61 | 0.85 | 0.64 | 0.78 | 0.73 | 0.84 | 0.85 | 0.87 | 0.78 |
| CNN-LSTM | 0.87 | 0.59 | 0.90 | 0.76 | 0.82 | 0.66 | 0.95 | 0.86 | 0.89 | 0.80 |
| ETRCNN | 0.84 | 0.66 | 0.87 | 0.76 | 0.57 | 0.89 | 0.80 | 0.89 | 0.90 | 0.80 |
| HDNN-TL | 0.92 | 0.63 | 0.86 | 0.67 | 0.81 | 0.75 | 0.86 | 0.87 | 0.91 | 0.81 |
| MSFFCNN-2 | 0.91 | 0.82 | 0.81 | 0.74 | 0.79 | 0.77 | 0.81 | 0.88 | 0.85 | 0.83 |
| **MSFFCNN-TL** | 0.91 | 0.89 | 0.87 | 0.86 | 0.89 | 0.85 | 0.83 | 0.87 | 0.92 | 0.88 |

**Figure 12:**
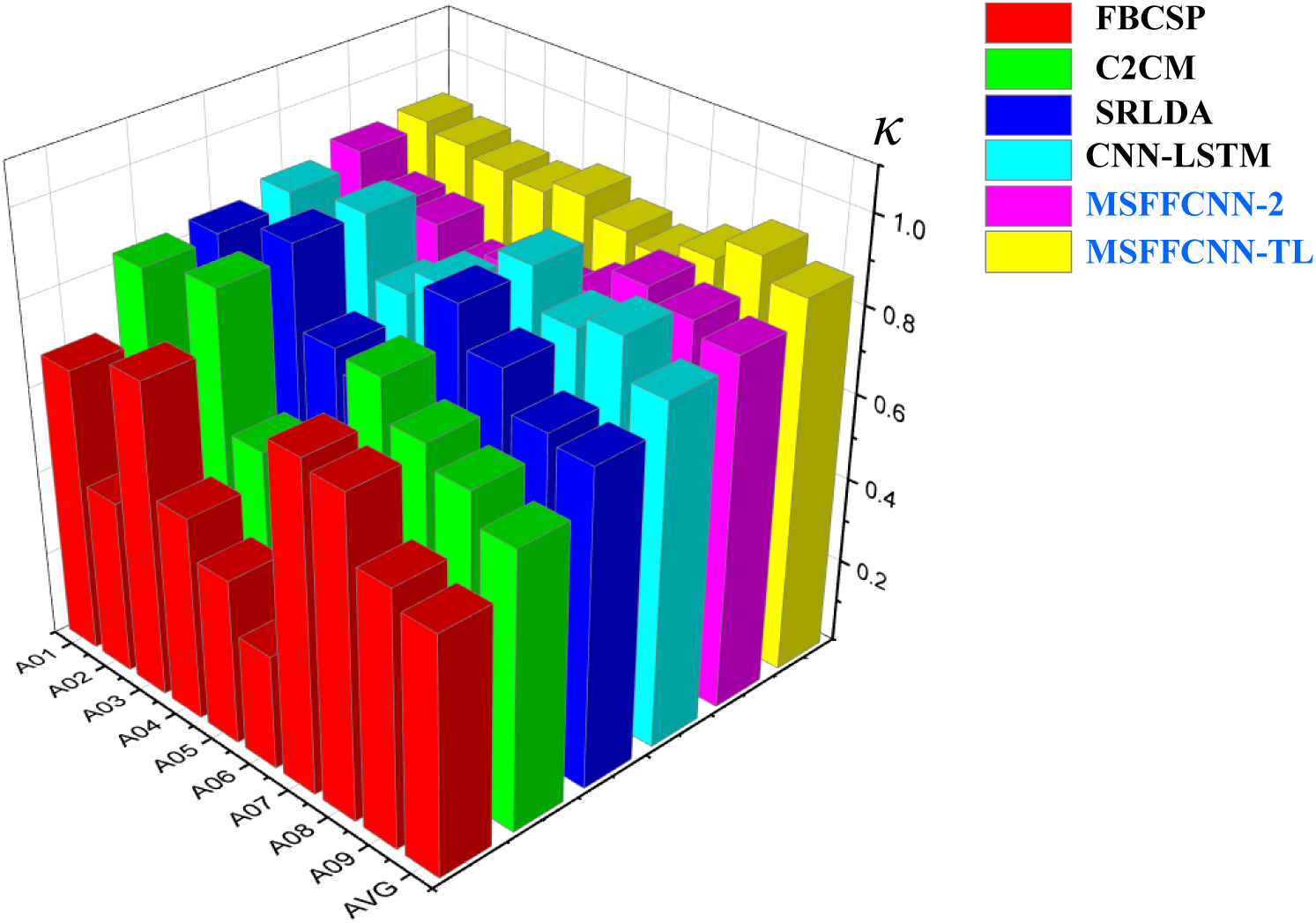
Comparison of *κ* across all subjects between state-of-the-art ML models and proposed MSFFCNN model.

### 6.4 Comparison with different state-of-the-art models

This section reports the accuracy comparison between proposed models and several state-of-the-art models such as support vector machine (SVM) [42], CW-CNN [42], transductive learning with covariate shift-detection (TLCSD) [72], C2CM [42], adaptive learning with CSD (ALCSD)[72], RBN *t*−SNE [69], densely feature fusion convolutional neural networks (DFFN)[73], subject specific multivariate empirical mode decomposition based filtering (SS-MEMDBF) [74], ETRCNN [71], Riemannian geometry-based FBCSP (RG-FBCSP) [75], and hybrid scale CNN (HS-CNN) [33] as outlined in Table 11. It can be seen that proposed MSFFCNN-2 and MSFFCNN-TL models attain the highest average *A*_*c*_ of 91.61% and 94.11% among all state-of-the-art models for BCI Competition IV 2a datasets. It is noteworthy to mention that MSFFCNN-TL improves the average classification accuracy of 14.12%, 8.51%, and 2.49% over the recent and advanced DL models SS-MEMDBF, RG-FBCSP, and HS-CNN, respectively as shown in Fig. 13. More specifically, MSFFCNN-TL demonstrated significant classification accuracy improvement of 7.70%, 3.67%, 3.03%, and 3.58% in subjects *A*02, *A*03, *A*05, and *A*09, respectively over MSFFCNN-2 which indicates the efficiency and robustness of the proposed subject-adaptive MSFFCNN-TL model.

**Table 11:**
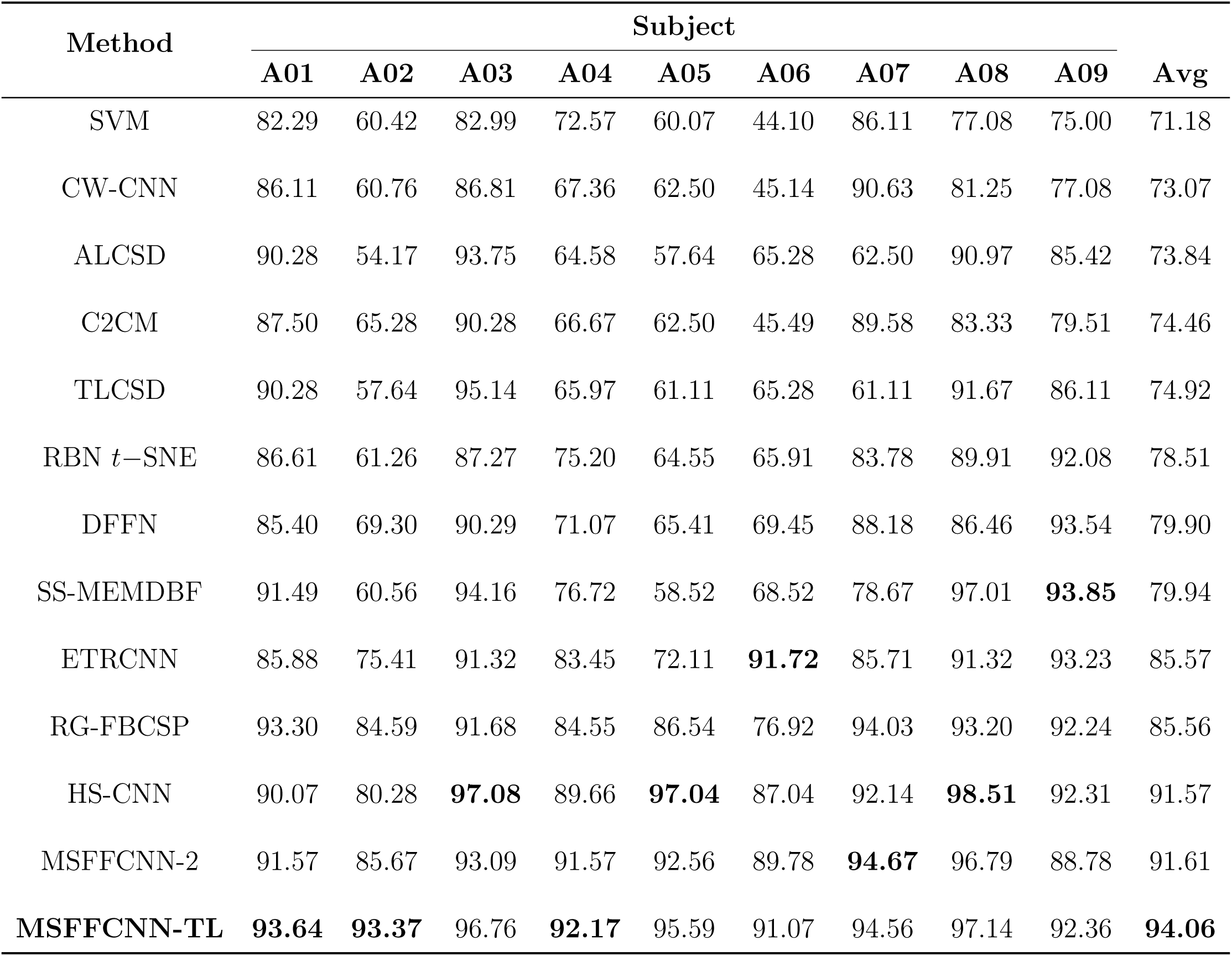
*A*_*c*_ for different subjects and average *A*_*c*_ (in %) obtained from existing state-of-the-art MI-BCI classification models.

**Figure 13:**
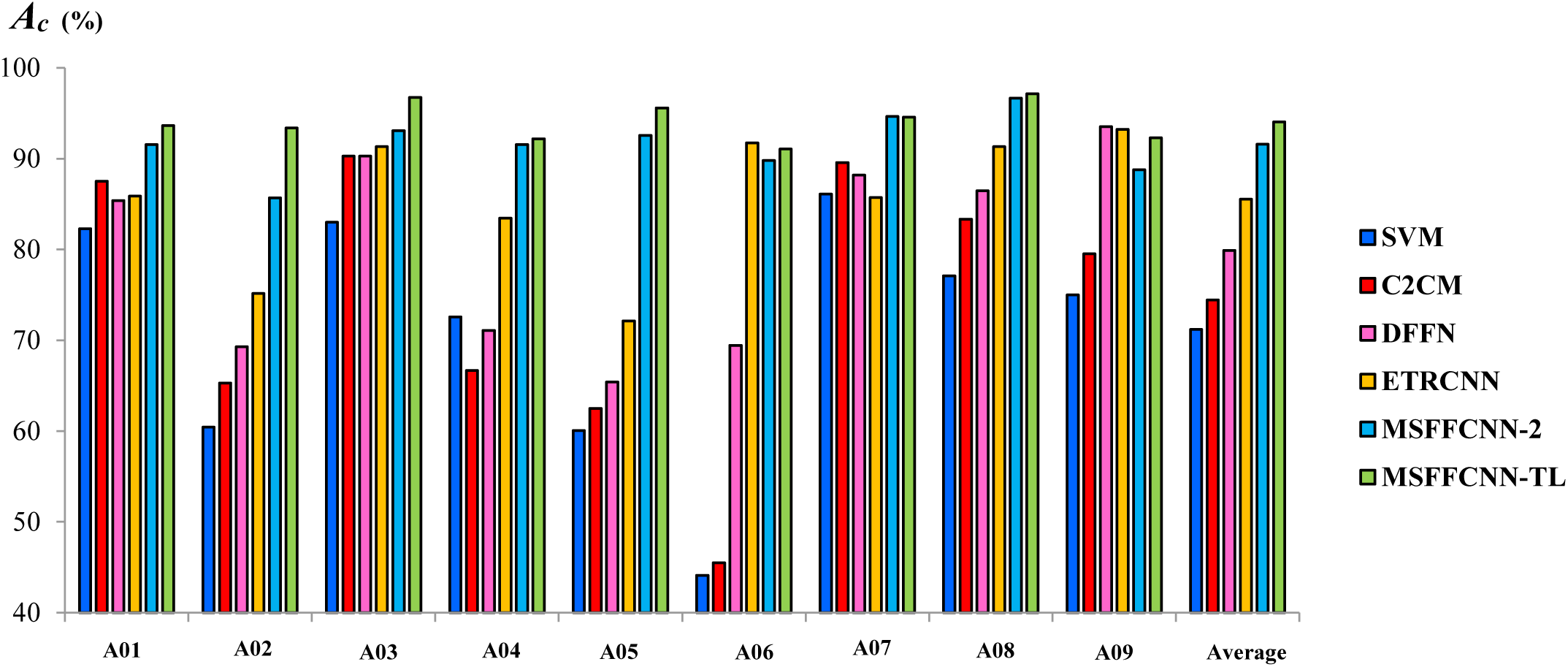
Comparison bar-chart of *A*_*c*_ for different subjects and average *A*_*c*_ (in %) between proposed MSFFCNN-2, MSFFCNN-TL and existing state-of-the-art MI-BCI models.

## 7. Discussion

The current research demonstrates the efficiency of the proposed MSFFCNN for extracting semantic features of EEG signals from multiple convolutional kernel scales for four different frequency bands to enhance the performance of the MI classifier. Incorporating OVR-FBCSP CNN further improves the accuracy of the classifier emphasizing the prospect of the current framework in adopting the distinguishable feature of MI-EEG signals. Furthermore, the current study illustrates the advantages of using an adaptive transfer learning-based CNN model over the subject-specific and subject-independent model in order to obtain better classification accuracy and performance. Due to inter-subject variability in MI-EEG data, the majority of the traditional ML methods in the MI-BCI system uses subject-specific data for better accuracy. However, to train a deep CNN network model containing a large number of trainable parameters requiring a significant amount of training data which is not suitable for relatively less subject-specific data limiting the accuracy of the classifier. In this regard, the current study shows the usefulness of subject-independent models which can be trained in a large number of inter-subject samples. In order to further increase the performance of the classifier, different adaptation techniques considering different learning rate scale factors, degree of adaption, and adaptation configurations have been explored to fine-tune the subject-independent model which provides significant improvement of accuracy of the classifier. The current study illustrates the importance of lowering learning rates which facilitate effective adaptation and improve classification accuracy. Among all adaptive models, it has been found out that for *θ* = 0.01 with a full degree of adaptation *ξ* = 1 and adaptation configuration *AC* − 2 provides the best average accuracy of 94.06% which is a 4.86% improvement over the subject-specific model. Thus, current work effectively addresses training and adaptation strategy in adaptive cross-subject transfer learning considering inter-subject variability for better performance. Future work will focus on further improving the classification accuracy by incorporating long short-term memory (LSTM) RNN architecture to extract temporal features and employ the proposed framework for classifying spatio-temporal multi-class MI subject classification for various BCI applications. Additionally, Maximum Mean Discrepancy (MMD) strategy [76] can be utilized to further regularize the adaptation of individual CNN layer. The proposed CNN framework can also be used in material informatics [77, 78, 79, 80, 81, 82]. Nevertheless, the proposed MSFFCNN model can be employed as a more reliable and robust MI-based real-time BCI applications such as robotic control [9, 10, 11], rehabilitation of neuromotor disorders [8], text entry speech communication [12, 13] etc.

## 8. Conclusion

Summarizing, in the current study, a transfer learning-based multi-scale featured fused CNN (MSFFCNN) framework has been presented for multi-class MI classification where the multiscale convolution block comprise of various convolutional kernel sizes can efficiently extract semantic features for different frequency bands *δ, θ, α*, and *β* in multiple scales. Various parametric exploration including the influence of learning rate scale factor and degree of adaptions on different adaptation configurations sheds light on effective and optimal adaptation strategies to maximize the performance of the proposed transfer learning model. The current study illustrates the importance of lowering learning rates which facilitate effective adaptation and improve classification accuracy. Among all adaptive models, it has been found out that for relatively low learning rate with a full degree of adaptation and adaptation configuration corresponds to a relatively larger number of numbers of adaptable network parameters provides the best average classification accuracy of 94.06% (±2.29%) which is a 4.86% improvement over the subject-specific model. The proposed framework requires less training data and computation time suitable for designing robust and efficient human-robot interaction. The present study effectively addresses the shortcoming of existing CNN-based EEG-MI classification models and significantly improves the classification accuracy.

